# *Mms19* promotes spindle microtubule assembly in neural stem cells through two distinct pathways

**DOI:** 10.1101/2020.06.08.139816

**Authors:** Rohan Chippalkatti, Boris Egger, Beat Suter

**Affiliations:** Institute of Cell Biology, University of Bern; Graduate School for Cellular and Biomedical Sciences, University of Bern; Department of Biology, University of Fribourg

**Keywords:** mitotic spindle, microtubule, Mms19, TACC, mitotic kinases, CAK complex

## Abstract

Mitotic divisions depend on the timely assembly and proper orientation of the mitotic spindle. Malfunctioning of these processes can considerably delay mitosis, thereby compromising tissue growth and homeostasis, and leading to chromosomal instability. Here we identified Mms19 as an important player in these processes as it promotes spindle and astral microtubule (MT) growth and consequently regulates spindle orientation and mitosis duration in *Drosophila* neural stem cells. Loss of functional *Mms19* drastically affects the growth and development of mitotic tissues in *Drosophila* larvae. We found that *Mms19* performs its mitotic activities through two different pathways. By stimulating the mitotic kinase cascade, it triggers the localization of the MT regulatory complex TACC/Msps (Transforming Acidic Coiled Coil/Minispindles, the homolog of human ch-TOG) to the centrosome. In addition, we present evidence that Mms19 stimulates MT stability and bundling by binding directly to MTs.

## Introduction

The fidelity of chromosome segregation during mitotic divisions depends on the proper regulation of the structure and dynamics of spindle microtubules. Numerous mitotic factors, including different mitotic kinases and microtubule (MT) associated proteins, meticulously regulate the formation, orientation, polymerization and catastrophe of spindle MTs to ensure that the chromatids are evenly delivered to the two products of the division process, the daughter cells. Incorrect regulation of Cdks and MTs can lead to chromosomal instability and to deleterious effects on the growth and development of somatic tissues.

*Mms19* was identified as a gene required for nucleotide excision repair (NER) (Prakash et al, 1979). The protein Mms19 can be found as part of the <u>C</u>ytoplasmic <u>I</u>ron-Sulfur <u>A</u>ssembly complex (CIA), which mediates the incorporation of iron-sulfur clusters into NER proteins such as Xpd (Gari et al, 2012; Stehling et al, 2012). However, a distinct, apparently DNA repair-independent, function of *Mms19* came to light when its knock-down caused numerous mitotic spindle abnormalities in human cells (Ito et al, 2010). Downregulation of *Mms19* in young *Drosophila* embryos also revealed spindle abnormalities and chromosome segregation defects, and these phenotypes were linked to the mitotic control pathway when results by Nag and co-workers revealed that Mms19 acts as a positive regulator of the <u>C</u>dk <u>A</u>ctivating <u>K</u>inase (CAK) activity (Nag et al, 2018). CAK has dual roles during the cell cycle where it activates the mitotic Cdk1 during mitosis, but is recruited by Xpd to form the holoTFIIH complex during interphase (Cameroni et al, 2010). The incorporation into TFIIH causes CAK to phosphorylate an entirely different set of substrates and allows CAK to perform its functions in transcription. Nag et al proposed that Mms19 binds to Xpd during mitosis and that this binding competes with the binding of Xpd to the TFIIH subunits. Binding to Xpd could thereby release CAK to activate its mitotic targets. In this model, reduced levels of Mms19 prevent the dissociation of the Xpd-CAK complex. As a consequence, the required levels of the mitotic CAK activity are not established. Indeed, overexpressing CAK complex components in the *Mms19* loss-of-function (*Mms19^P^*) background rescued the mitotic defects to a large degree (Nag et al, 2018).

The remarkable findings by Nag et al uncovered a novel mitotic pathway for *Mms19*. But this study mostly focused on the young *Drosophila* embryo which is unusual in that all somatic nuclei share a common cytoplasm in which the mitotic divisions take place. Furthermore, their cell cycle consists of only S and M phases, without intervening G phases. In this situation with the shared cytoplasm, the absence of Mms19 often causes microtubules emanating from one spindle pole to contact the chromosomes of a neighboring nucleus. It is therefore difficult to extrapolate these findings to mononuclear diploid cells that are isolated from their neighbors by plasma membranes and have a full cell cycle. Furthermore, even though the spindle abnormalities observed in the absence of Mms19 could be linked to compromised CAK activity, the pathway acting downstream of CAK is not clearly understood. Finally, overexpression of the CAK complex brought about only a partial rescue of the *Mms19^P^* defects, pointing towards additional, possibly CAK-independent spindle regulatory roles of *Mms19*.

The objective of this study was thus to investigate the mitotic function of *Mms19* in normal diploid cells with the goal of dissecting the precise pathway through which *Mms19* acts to regulate mitotic spindle assembly and cell cycle progression. Because *Mms19^P^* larvae lack imaginal discs, we chose the larval brain neuroblasts (NBs) as a model to analyze the mitotic roles of *Mms19* in cells with a full cycle. We identified several novel *Mms19* phenotypes, which allowed us to pinpoint steps in the pathways that require *Mms19* activity. We found that *Mms19* is required in NBs for timely progression through the cell cycle and consequently for establishing normal cell numbers in the NB lineage. *Mms19* is also required for the growth and assembly of spindle and astral microtubules (MTs). Our results connect these defects to the mis-localization of the microtubule regulator TACC, suggesting that TACC is a downstream target of the mitotic kinase cascade through CAK-Cdk1-Aurora kinases. Additionally, and apparently unrelated to the CAK-Cdk1 axis, we also identified a direct interaction *in vitro* between Mms19 and microtubules and found that Mms19 promotes microtubule stability and bundle formation.

## Results

### *Mms19^P^* brains show a microcephaly phenotype

Normal *Drosophila* larvae spend on average 5-7 days at 25°C to pass through the larval stages. In contrast, *Mms19^P^* larvae take not only at least 8-10 days to reach the size of outgrown WT third instar larvae, but they also spend a total of around 15 days in the 3^rd^ larval instar stage before they die. These larvae display a typical mitotic phenotype without recognizable imaginal disc tissues (Nag et al, 2018). Even though outgrown *Mms19^P^* larvae lack imaginal discs, their brain is still present, although it is much smaller, displaying a microcephaly phenotype (Fig 1A-C). Additionally, the optic lobe (OL) appears deformed or underdeveloped. Compared to the wild type, the total volume of the *Mms19^P^* brains and the volumes of the CBs and OLs, too, are drastically reduced (Fig 1E-G). Surprisingly, however, the number of central brain (CB) NBs per brain lobe did not significantly change between the controls and the *Mms19^P^* brains (Fig 1D). This implies that the *Mms19^P^* NBs have been properly determined and are present, but probably divide slowly, thereby contributing fewer Ganglion Mother Cells (GMCs) and neurons to the CB and resulting in reduced brain size. We expressed wild-type Mms19 fused to eGFP (Mms19::eGFP) under the control of the endogenous *Mms19* promoter in the *Mms19^P^* background to test whether the observed phenotype was indeed caused by reduced *Mms19* activity. Mms19::eGFP fully rescued the brain morphology, the volumes of the whole brain lobe as well as the CB and OL sizes (Fig 1A-G). This not only confirmed that the defects observed in the mutant are indeed due to the absence of *Mms19* activity and not due to a second site mutation on this chromosome, it also showed that the Mms19::eGFP fusion protein is functional.

**Figure 1.**
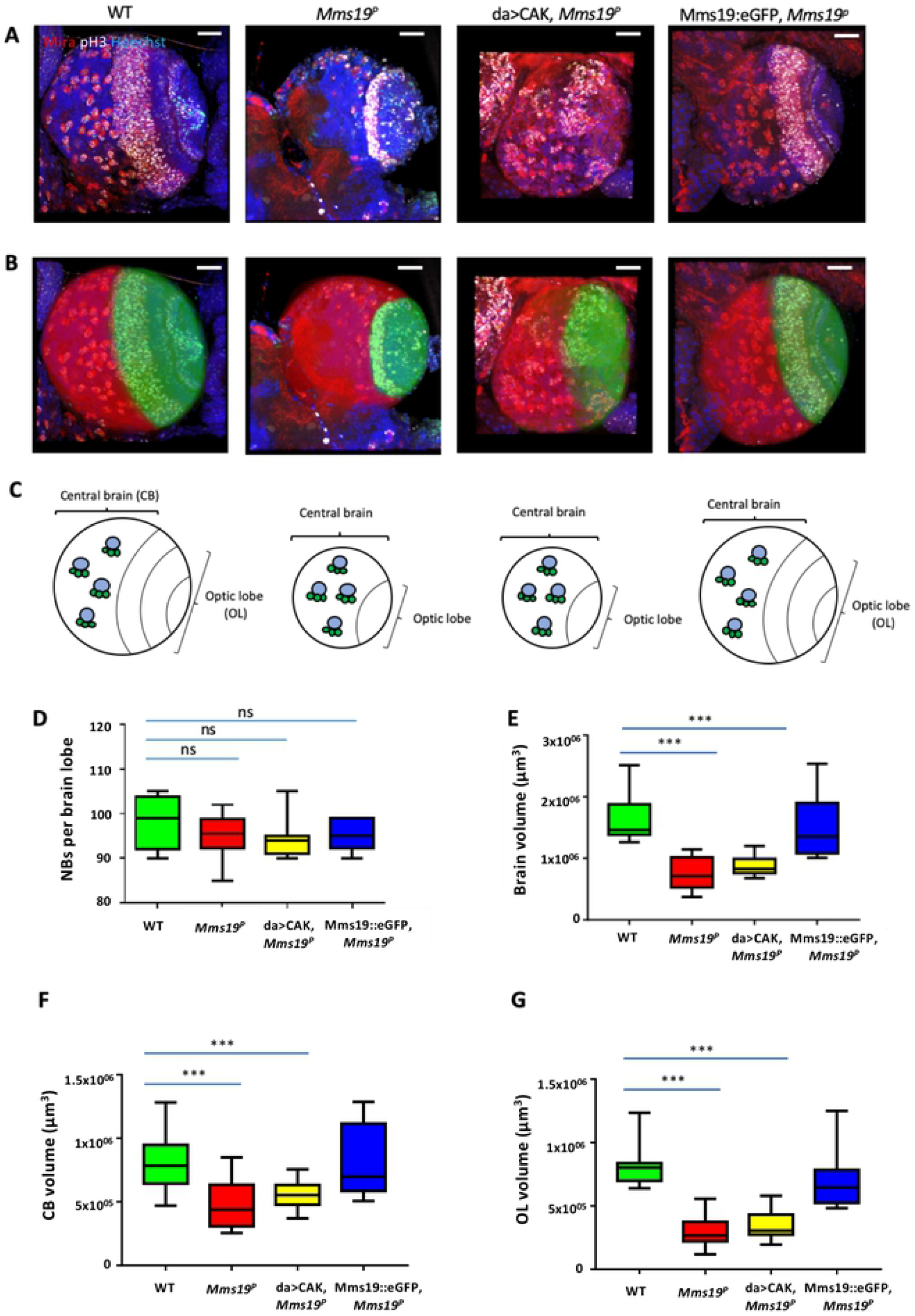
*Mms19^P^* brains display a microcephaly (small brain) phenotype. 3^rd^ instar larval brain NBs were visualized by staining for Miranda (red, cytoplasmic) and pH3 (white, nuclear); DNA (Hoechst 33342 dye) is shown in blue. (A)-(C) A WT brain lobe can be subdivided into the OL (shaded green) and the CB (shaded red). *Mms19^P^* brains appear smaller, with a diminished OL. Overexpressing the CAK complex components driven by daughterless-Gal4 (da>CAK) in the *Mms19^P^* background did not rescue the phenotype. Mms19::eGFP expressed in the *Mms19^P^* background rescued the *Mms19^P^* phenotype. Although the number of NBs per brain lobe is similar across all genotypes (D), the volume of the brain lobe is significantly reduced in the *Mms19^P^* and da>CAK, *Mms19^P^* brains (E). Furthermore, segmentation and volume measurement of the OL and CB revealed a significant reduction in *Mms19^P^* and da>CAK, *Mms19^P^* brains. The brain volume and morphology were restored to WT levels when Mms19::eGFP was expressed in *Mms19^P^* brains (E)-(G). n=15 (15 brains, 30 lobes). Statistical significance (SS) was determined by the Kruskal-Wallis test. Multiple columns were compared using Dunn’s posttest, ***(*P*<0.001), scale=25μm.

A mitotic activity of *Mms19* was described to promote the Cdk-activating kinase activity of the Cdk7/CycH/Mat1 complex (CAK complex; Nag et al, 2018). The responsible mechanism appears to be a competitive binding of Mms19 to Xpd, which would otherwise recruit CAK to TFIIH, where it assumes a different substrate specificity and is unable to activate the M-Cdk Cdk1. The key result that lead to this model was that the lack of imaginal discs caused by the absence of *Mms19* activity could be rescued considerably by expressing the CAK complex components under the control of the <u>U</u>pstream <u>A</u>ctivating <u>S</u>equence (UAS) enhancer, using the *daughterless*(*da*)-Gal4 driver in the *Mms19^P^* background (da>CAK, *Mms19^P^*) (Nag et al, 2018). We therefore tested whether it was possible to rescue the small brain phenotype by the identical strategy. This was not the case, as both CBs and OLs remained smaller (Fig 1A-G). In da>CAK, *Mms19^P^* brains, the number of CB NBs was similar as in the wild type (Fig 1D), even though the overall brain volume was reduced. These observations indicated that the *Mms19^P^* CB NBs probably did not proliferate enough to produce the normal amount of neuronal tissue, and that this defect might not only be caused by insufficient CAK activity. This result therefore points to the possibility that Mms19 acts through two different pathways to achieve normal organ size.

### Higher fraction of *Mms19^P^* NBs in mitosis

In order to better understand the microcephaly phenotype, we performed 5-Ethynyl-2’-deoxyuridine (EdU) incorporation assays coupled with pH3 staining and examined defects in cell cycle progression in the NBs. Based on pH3 and EdU staining, the cells can be allocated to one of the following phases: only EdU = S, i.e. cells in S phase; pH3 without EdU = M, i.e. cells undergoing mitosis; both EdU and pH3 =G2/M, i.e. cells transiting from S to G2/M phase; and neither EdU nor pH3 (= G1/G0, i.e. Gap phase) (Fig S1A). NBs were marked with antibodies against Miranda (Mira) and the number of cells were counted in each class. The results are represented as a percentage of the total number of cells per brain lobe. We observed that about half the cells were not labelled (i.e. G1/G0 cells: 47-56%; Fig S1B) and the relative differences between the genotypes were small. However, lack of *Mms19* caused a clear increase in the fraction of cells in M phase (35% compared to 24% in the wild type; Fig S1E). This result could either mean that more cells undergo divisions or that the *Mms19^P^* NBs are either trapped in M phase or proceed more slowly through it.

Interestingly, the small class of cells that are EdU^+^ and pH3^+^ showed the highest relative increase in frequency when CAK was overexpressed (Fig S1D; 6.5% vs 3.2% for *Mms19^P^*). Double positive cells have gone through S phase and were in M-phase at the time of fixation. The fact that overexpression of CAK leads to a higher frequency of double positive cells is consistent with the idea that these cells are progressing through the cell cycle more rapidly and/or that they enter M-phase prematurely.

### NBs depend on *Mms19* for timely and coordinated spindle assembly and spindle orientation

To study how *Mms19* contributes to spindle assembly and progression through mitosis, we utilized a transgene that expresses EB1::GFP, a MT plus end binding and tracking protein that labels growing MT ends (Zhu et al, 2009). Live imaging of NBs expressing EB1::GFP revealed the dynamics of the formation of the mitotic spindle and allowed us to estimate the duration of mitosis, which we defined as the period from the onset of the <u>N</u>uclear <u>E</u>nvelope <u>B</u>reak-<u>D</u>own (NEBD) until the end of cytokinesis. NEBD onset was determined by the appearance of the GFP signal in the nuclear region, which lacks a GFP signal until NEBD. NBs expressing EB1::GFP in the wild-type background finished cytokinesis approximately 10 min after NEBD (Fig 2A, mov 01). On the other hand, NBs expressing EB1::GFP in an *Mms19^P^* background reached cytokinesis only around 20 min after NEBD (Fig 2B, mov 02).

**Figure 2.**
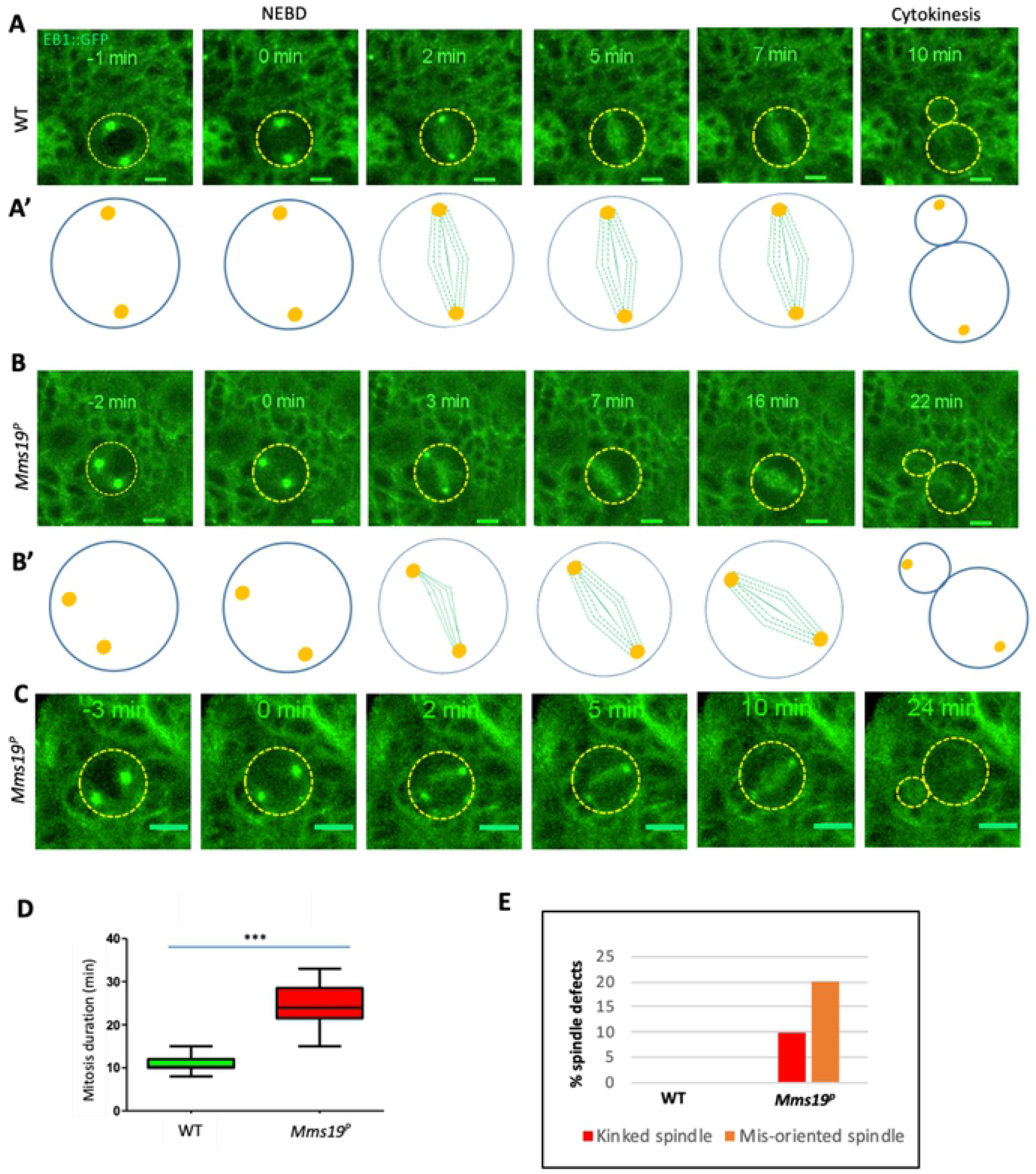
NBs depend on *Mms19* for timely and coordinated spindle assembly and spindle orientation. (A) WT NBs typically finish cytokinesis 10 min after the onset of nuclear envelope breakdown (NEBD). (B) *Mms19^P^* NBs require on average 20 min to reach cytokinesis after NEBD. (B, B’) In around 10% the mutant cells, the spindle poles are not fully separated at NEBD and the spindle initially appears kinked. In around 20% of *Mms19^P^* NBs, the spindles change their orientation throughout the course of mitosis. (C) In this particular mutant case, MTs are initially seen to emanate only from a single pole, but subsequently, the other pole also nucleates MTs. The quantitative assessment of the duration of WT and *Mms19^P^* mitoses is compared in (D). SS was determined by an unpaired t-test (\*\*\**P*<0.001); Scale=5μm; n=30 NBs. Kinked spindles and mis-oriented spindles are quantified in (E).

Interestingly, in some of the EB1::GFP expressing *Mms19^P^* NBs, the spindle formation started before the two centrosomes had finished migrating to the opposite sides of the nucleus (Fig 2B, mov 03). As a result, at 3 min, a kinked spindle was observed, which eventually straightened out at 5 mins. Around 10% kinked spindles were found in *Mms19^P^* NBs. In one case of a *Mms19^P^* NB spindle, only one centrosome started nucleating MTs (Fig 2C). The spindle remained monopolar until 6 min post NEBD and only then, a bipolar spindle became apparent. Further, around 20% of the spindles in *Mms19^P^* NBs changed their orientation during the course of mitosis (Fig 2B, B’, mov 03), while all wild type spindles examined remained firmly anchored at the cortex and did not change their orientation (Fig 2A, A’).

*Drosophila* larval NBs are characterized by a defined apical-basal polarity (Homem et al, 2012) where the atypical Protein Kinase C (aPKC)-Bazooka-Pins complex localizes to the apical cortex while Miranda localizes to the basal cortex (Fig S4A). As differentiation of the GMC relies on the inheritance of the Mira-bound cargo, the orientation of the mitotic spindle is tightly coupled to this apical basal polarity axis (Schuldt et al, 1998; Rolls et al, 2003). We therefore further examined the spindle orientation defects in fixed cells labelled with the polarity marker Miranda (Mira), which localizes to the basal cortex of the NB, forming a crescent-like pattern. We found that most of the WT spindles form at an angle of 10 degrees or lower relative to Mira. On the other hand, *Mms19^P^* NBs more frequently failed to align their spindles within 10 degrees (Fig S4C-D) and the largest tilt that we observed was around 60°. However, a marked effect on cell fate determination was observed previously only when the spindle angle was close to 90° (Lee et al, 2006; Cabernard and Doe, 2009). In this case spindles oriented at around 90° led to both the daughter cells assuming the NB identity, thus increasing NB numbers per brain lobe. Consistent with this spindle orientation defect being insufficient to cause major NB amplification, we did not see a significant difference in the NB number per brain lobe between wild-type *Mms19^P^* (Fig 1D). Therefore, the spindle orientation defect due to lack of *Mms19* does not appear to lead to differentiation problems, but impedes efficient mitosis.

We conclude that in the absence of *Mms19*, spindle formation is not properly coordinated with cell cycle progression and that defects in centrosome migration and in spindle assembly and orientation contribute to the delay of the mitosis in *Mms19^P^* NBs.

### *Mms19* is cell autonomously required to maintain normal cell numbers

In the experiments with the *Mms19^P^* mutants, Mms19 was absent from all larval cells. In this situation, the mitotic delay could be due to a systemic effect or due to the lack of a cell autonomous activity of Mms19. To test whether *Mms19* is specifically required in the mitotic cells for the timely progression through mitosis, we generated *Mms19^P^* mosaic NB clones in an otherwise wild-type background. Mosaic clones were induced in NBs 24hrs after larval hatching (ALH) and the expansion of these clones was analyzed by dissecting the brains in mid-third instar larvae 72 hrs ALH. For this experiment, we focused on the type I NBs on the ventral side. On average, 45 cells per clone were counted in control clones, but only around 30-35 cells in *Mms19^p^* clones (*P*<0.05; Fig S2 A-C), indicating that the loss of *Mms19* activity hinders the establishment of normal cell numbers in the NB lineage. This observation reaffirms our conclusion that absence of *Mms19* delays mitosis in NBs and that this mitotic delay probably causes more *Mms19^P^* NBs to linger in M-phase (Fig S1E).

### *Mms19* is required to form spindles of a normal length and density

*Mms19^P^* spindles are generally shorter than the wild-type ones (Fig 3A-E). In normal NB spindles, the centrosomes were anchored close to the cell cortex, with spindle MTs emanating from them and extending to the chromosomes (Fig 3A, A’). But in around a quarter of the mutant cells, even though the chromosomes were aligned at the metaphase plate, the centrosomes were connected to the cell cortex and were only a short distance away from the metaphase plate (Fig 3B, B’). In order to quantify this defect, we measured the length of the spindles across all genotypes and found that the spindles in *Mms19^P^* and da>CAK, *Mms19^P^* were significantly shorter than the wild-type control spindles (Fig 3C). As the NBs were also considerably smaller in *Mms19^P^* and da>CAK, *Mms19^P^* brains (Fig 3D), we additionally displayed the spindle length relative to the cell diameter (Fig 3E). A ratio closer to 1 indicates that the centrosomes were anchored close to the cell cortex, as in a healthy spindle. On the other hand, if the ratio was equal to or lower than 0.6, the spindle was defined as a ‘short spindle’ because the centrosomes were further inside the cell. We found a significant reduction in this ratio in *Mms19^P^* and da>CAK, *Mms19^P^* NB spindles (Fig 3E). *Mms19^P^* NBs contained around 20% short spindles while this number went up to 30% for da>CAK, *Mms19^P^* (Fig 3F). This data indicates that the short spindle phenotype was not rescued, but actually worsened upon overexpression of CAK. On the other hand, expression of Mms19::eGFP in the *Mms19^P^* background rescued the short spindle phenotype and only 3% of the NBs displayed it.

**Figure 3.**
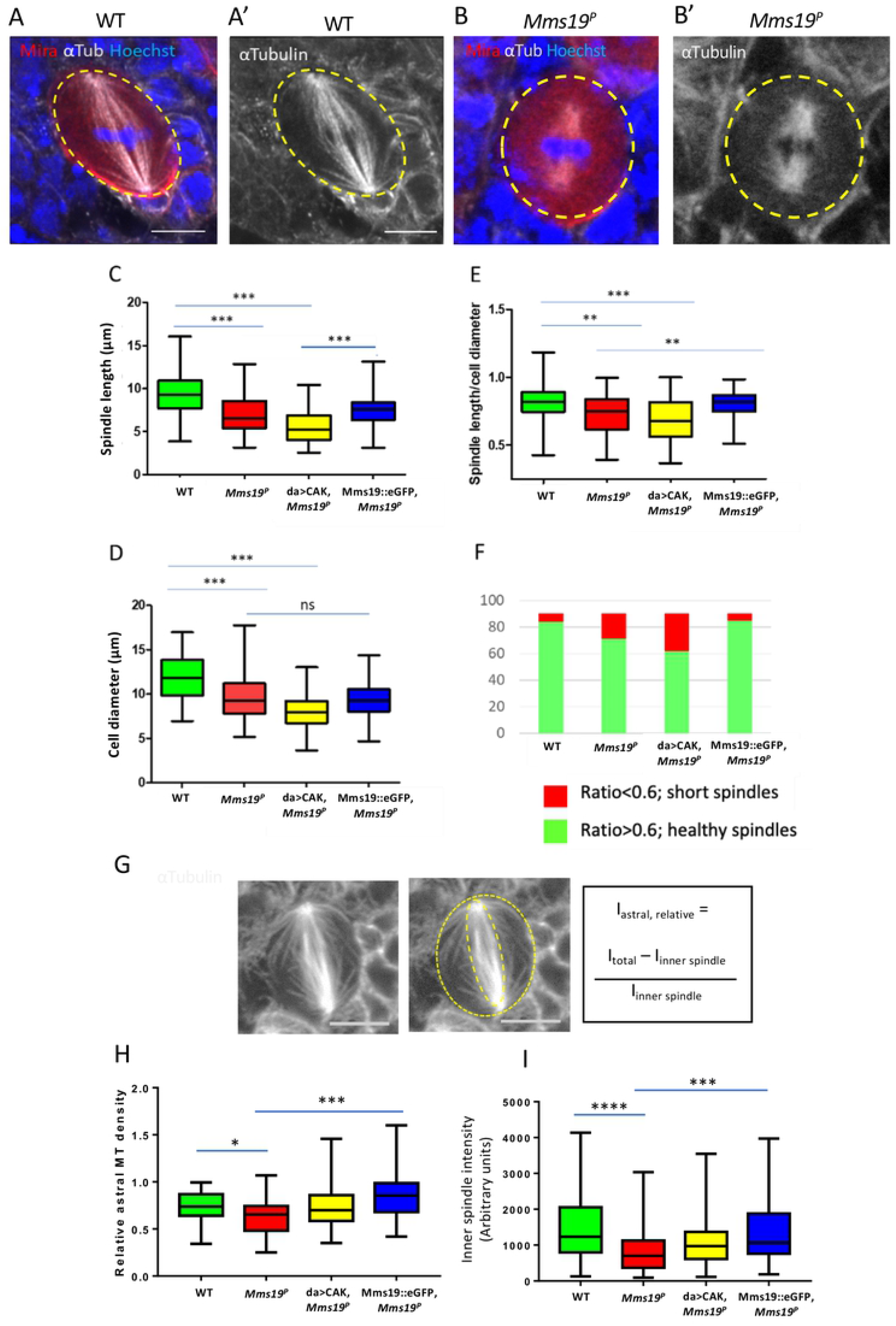
*Mms19* is required to form a spindle of normal length and density. (A-A’) Z projection of a typical bipolar spindle in a WT NB with the spindle poles anchored to the cell cortex. (B-B’) short spindle found in *Mms19^P^* NBs. the spindle is abnormally short and the spindle poles are detached from the cortex. (C) WT spindles are significantly longer than *Mms19^P^* and da>CAK, *Mms19^P^* spindles. (D) Graph comparing cell diameters between the different genotypes. WT NBs are bigger than *Mms19^P^* and da>CAK, *Mms19^P^* cells. (E) In order to quantify the short spindles, I normalized the spindle lengths to the cell diameters. If this ratio was equal to or less than 0.6, I considered these as ‘short spindles’. (F) Graph comparing the percentage of ‘short spindles’ across different genotypes. n= 90 NBs. Relative density of astral MTs was quantified and compared across all genotypes. Maximum intensity projections of mitotic NBs were obtained and the relative density of astral MTs was quantified as shown in (G). (H) Astral MT density and (I) Inner spindle fluorescent intensity was measured in fixed NBs immunostained for αTubulin and compared across all the genotypes n= 60 NBs. SS was calculated using Kruskal-Wallis test, columns were compared using Dunn’s post test (\*\*\*\**P*<0.0001, \*\*\**P*<0.001; \*\**P*<0.01, \**P*<0.05), scale=5μm.

To determine the effect of *Mms19* on MT formation and stability, we measured astral as well as inner spindle MT density. For astral MT quantification, we used the method described by Yang and co-workers (Yang et al, 2014; Fig 3G) and found a significant reduction in the *Mms19^P^* NBs as compared to wild type (Fig 3H-I). This phenotype appeared to be slightly rescued by CAK complex overexpression and was fully rescued by expressing Mms19::eGFP in the *Mms19^P^* background. Reduced astral MT stability could be linked to the spindle positioning and orientation defects described previously (Fig 2B, 2E, S4B-D) as astral MTs were shown to contact the cell cortex and regulate spindle positioning (Pearson et al, 2004). *Mms19* is thus necessary for the formation of fully assembled spindles and astral MTs.

### *Mms19* assists MT polymerization *in vivo*

To assess whether the short spindles could result from a defect in MT growth *in vivo*, we studied NBs expressing EB1::GFP. Live imaging of EB1::GFP revealed the path of elongation of single MTs, akin to that of a comet, and the movement speed of the GFP signal reflects the growth speed of the MT plus ends. Wild-type and *Mms19^P^* larval brains expressing EB1::GFP were dissected and live imaging was performed. The speed of the EB1::GFP particles was then tracked manually using ImageJ. The measurements indicated that the spindle MTs of wild-type NBs polymerized at a rate that is significantly higher than the rate observed in *Mms19^P^* NBs (Fig 4A, A’, WT-mov 04, *Mms19^P^*-mov 05). To assess whether the MT assembly function of *Mms19* is restricted to the mitotic state, we also tested whether the absence of *Mms19* also affects MT polymerization in the post mitotic glia cells. Measuring EB1 comet speeds in glia cells showed a decrease in MT growth for *Mms19^P^* glia as compared to wild-type glia (Fig 4B, B’, WT-mov 06, *Mms19^P^*-mov 07). These results established that *Mms19* assists MT polymerization and growth *in vivo* in mitotic NBs and in postmitotic glia.

**Figure 4.**
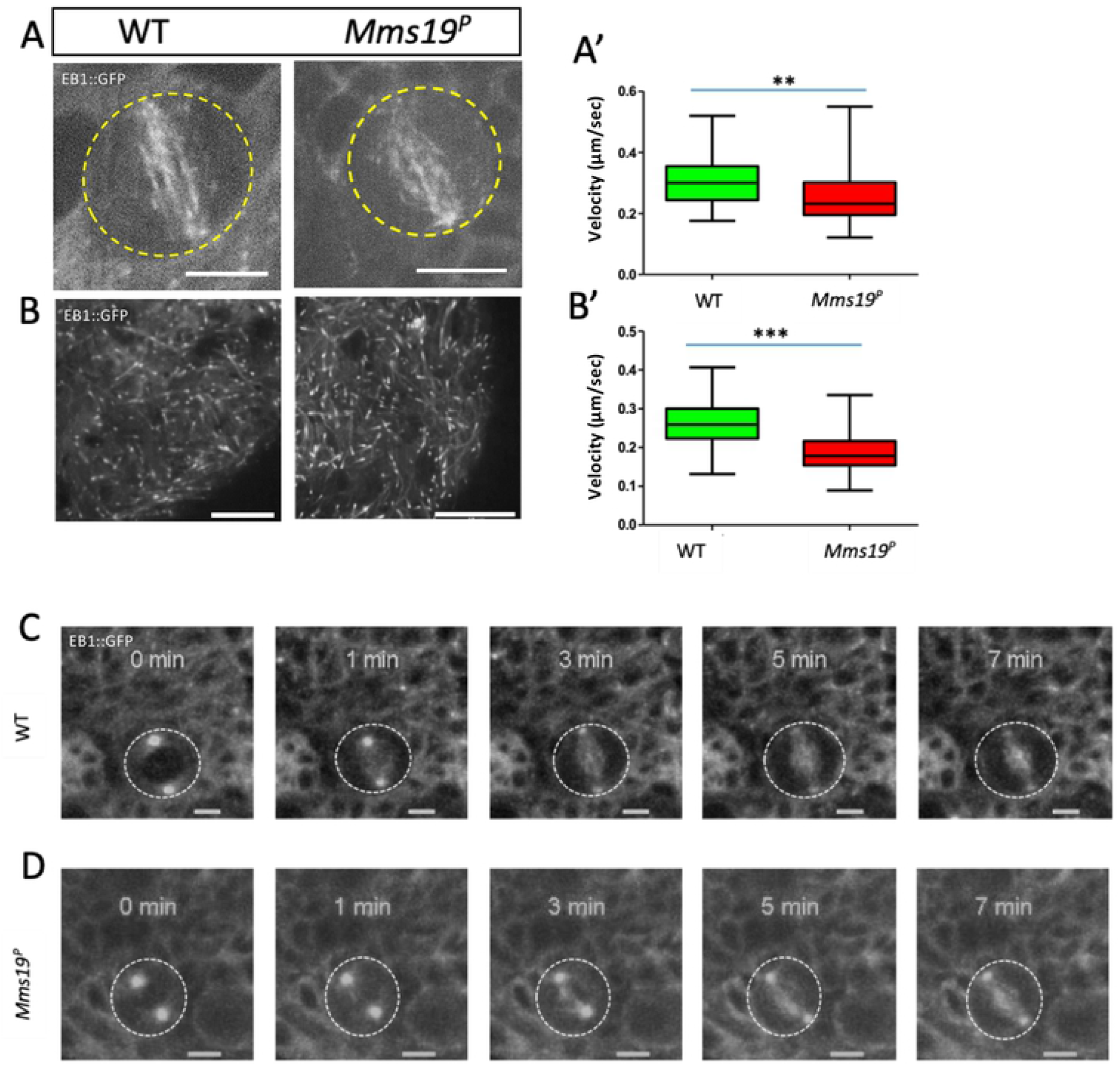
*Mms19* assists MT polymerization *in vivo*. (A) WT and *Mms19^P^* NBs expressing EB1::GFP were imaged live and time-lapse movies were acquired to calculate the velocity of EB1::GFP labelled MT ‘plus’ ends. The velocities represented in (A’) show reduced MT growth velocity in *Mms19^P^* NB spindles. n=11 cells, 5 particles analyzed from each cell (B-B’) EB1::GFP velocities were compared across WT and *Mms19^P^* in the glia. SS for A’ and B’ was calculated using unpaired t-test, (\*\**P*<0.01) (\*\*\**P*<0.001), scale bar=5μm. n=15 brains, 10 particles analyzed from each brain (C) In a WT NB, a spindle is seen to be fully assembled approximately 3 minutes after NEBD. (D) On the other hand, in most *Mms19^P^* NBs, spindle assembly seemed to be delayed.

Perhaps due to reduced MT growth, spindle assembly is also delayed in *Mms19^P^* brain NBs. Approximately 3 minutes after the onset of NEBD, a spindle fully assembled in wild-type NBs (Fig 4C). In contrast, in most *Mms19^P^* NBs, 3 minutes after the onset of NEBD the ‘spindles’ appear to be short and MT density seems much lower than in the wild type (Fig 4D). In the presented case, it took 7 minutes from the start of NEBD until a fully assembled spindle became visible (Fig 4D).

### *Mms19* is required for spindle re-polymerization

The short spindles observed in *Mms19^P^* NBs might be caused by impaired MT polymerization. To test whether Mms19 modulates MT polymerization dynamics, we utilized the *in vivo* MT polymerization assay described by Gallaud and co-workers (Gallaud et al, 2014). With this procedure, the NB spindles were completely depolymerized after incubation on ice for 30 min (Fig 5A). Wild-type NB spindles then regained their standard size and morphology within 90 sec after shifting them back to 25°C (Fig 5A). In *Mms19^P^* NBs, however, the spindles failed to re-polymerize to the normal shape after 90 sec. Instead, they remained abnormally short (Figs 5B, 5E). Interestingly, whereas the Mms19::eGFP fusion protein was able to rescue this phenotype, CAK overexpression was unable to do so (Fig 5C-E). To validate this short spindle phenotype, we also calculated the spindle sizes relative to the cell diameter after 90 sec incubation at 25°C (Fig 5F). Even though *Mms19^P^* cells are smaller, their normalized spindle size was still significantly smaller than the wild-type one. When CAK was overexpressed in *Mms19^P^*NBs, the spindle became slightly longer than in *Mms19^P^*, but it still remained shorter than in the *Mms19^P^* mutant rescued with the wild-type Mms19::eGFP transgene (Fig 5F). This result shows that *Mms19* has an important role in promoting spindle polymerization and mainly through a CAK independent process.

**Figure 5.**
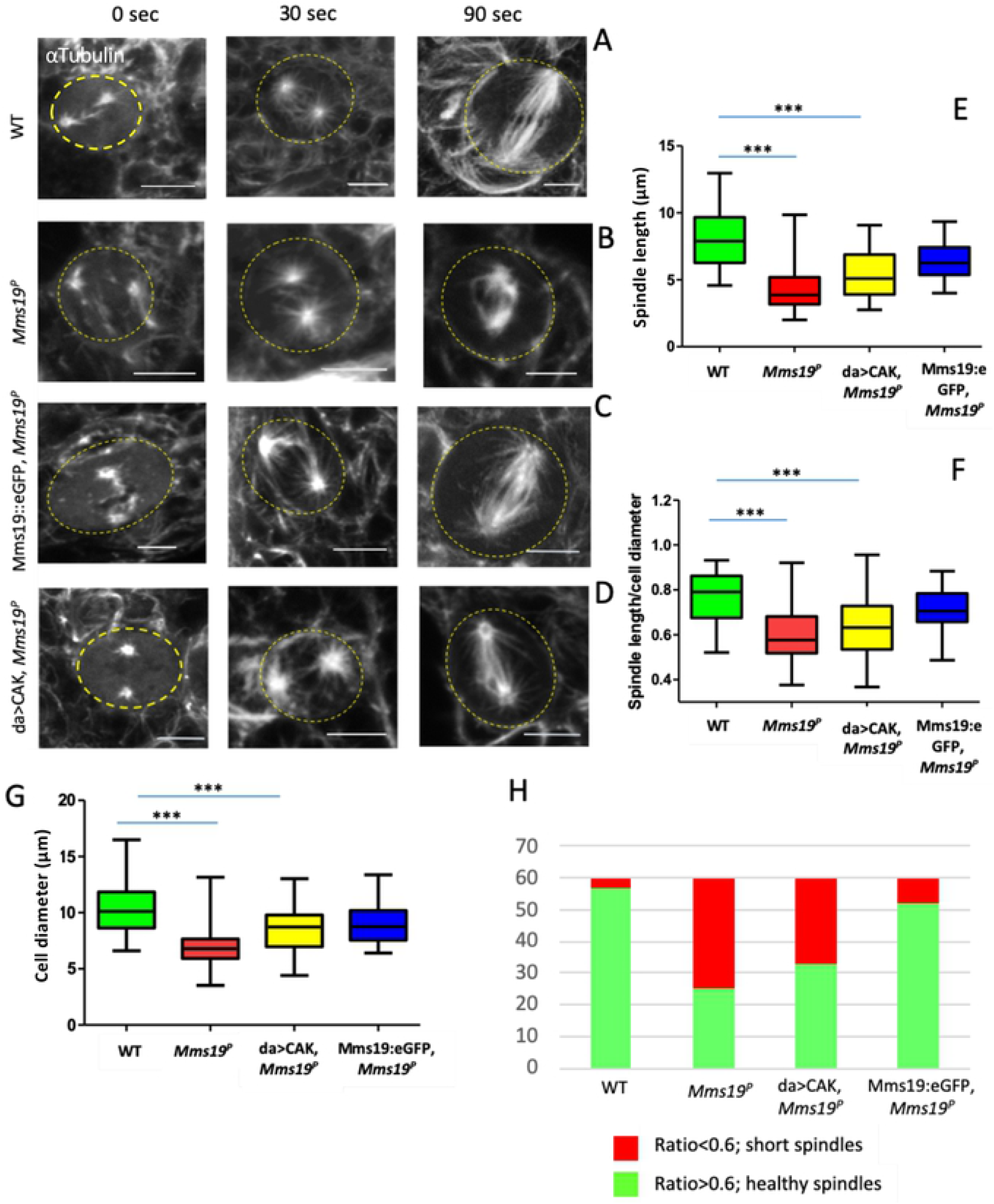
*Mms19* is required for spindle re-polymerization. After cold treatment, spindle re-assembly was analyzed at 0 sec (i.e. immediately after cold treatment; left panel), 30 sec (central panel) and 90 sec (right panel) after shifting them to 25°C. At 0 sec in the WT (A), only the centrosomes were visible but after incubation at 30 sec a few fibers nucleated from the centrosomes. At 90 sec, the spindle regained its normal shape. In *Mms19^P^* NBs (B), spindles did not regain the normal shape after 90 sec. Additionally, the microtubule density was reduced in the stunted spindles. Expression of Mms19::eGFP in the mutant background (C), rescued spindle reformation, but overexpression of CAK (D) rescued only partially and the length and density of MT still appeared reduced. (E) Chart comparing length of spindles after 90 sec incubation at 25°C. (F) The box plot (G) shows the spindle length relative to the cell diameter, thus eliminating variations due to varying cell sizes. SS was calculated by Kruskal-Wallis test, columns were compared by Dunn’s posttest (\*\*\**P*<0.001), scale=5μm, n=60 (F) The graph compares the percentage of ‘short spindles’ across different genotypes.

### Centrosomal localization of the MT regulator TACC depends on *Mms19*

Transforming Acidic Coiled-Coil (TACC), a downstream target of Aurora A kinase, is a critical regulator of centrosomal MTs. During mitosis, Aurora A phosphorylates TACC and stimulates its localization on the centrosomes, where TACC further recruits mini-spindles (Msps). Loss-of-function mutations in either *Aurora A, TACC* or *Msps* show drastic abnormalities in astral and spindle MTs (Giet et al, 2002; Bird and Hyman, 2008). As *Mms19^P^* NBs display defects in centrosomal separation, spindle length and astral MTs, we examined whether Aurora A and TACC might be involved in the same process as Mms19 and if the absence of functional *Mms19* impedes *TACC* function. We therefore stained wild-type and *Mms19^P^* NBs for TACC and observed a strong signal at the wild-type centrosomes (Fig 6A). On the other hand, in >50% of *Mms19^P^* NBs, TACC did not show any enrichment on the centrosomes (Fig 6B, D). Similar results were also obtained with its interactor Msps (Fig S6). As TACC acts downstream of Aurora A kinase, and Aurora itself acts downstream of Cdk1 (Van Horn et al, 2010), this defect might be caused by insufficient CAK activity in *Mms19* mutants (Nag et al, 2018). We therefore tested this hypothesis by over-expressing the three CAK components in the *Mms19^P^* background (da>CAK, *Mms19^P^*). Indeed, upon CAK overexpression, the fraction of spindles displaying centrosomal TACC almost reached wild-type levels (Fig 6C, D). To quantify the enrichment of TACC on centrosomes, we compared the fluorescent intensity of centrosomal TACC to the TACC fluorescent intensity on the spindle. This ratio decreased for *Mms19^P^* NBs, but was rescued in da>CAK, *Mms19^P^* NBs (Fig 6E). It was also reported that *Aurora A* loss-of-function can cause centrosome fragmentation (Giet et al, 2002). Co-staining *Mms19^P^* NBs with antibodies against the centrosomal protein g-Tubulin showed that even in cases with apparent TACC mis-localization, centrosomal g-Tubulin was still present, indicating that TACC mislocalization is not caused by centrosome fragmentation. These findings indicate that centrosomal localization of TACC and its stimulation of astral and spindle MT stability or growth is at least partially dependent on *Mms19* activity.

**Figure 6.**
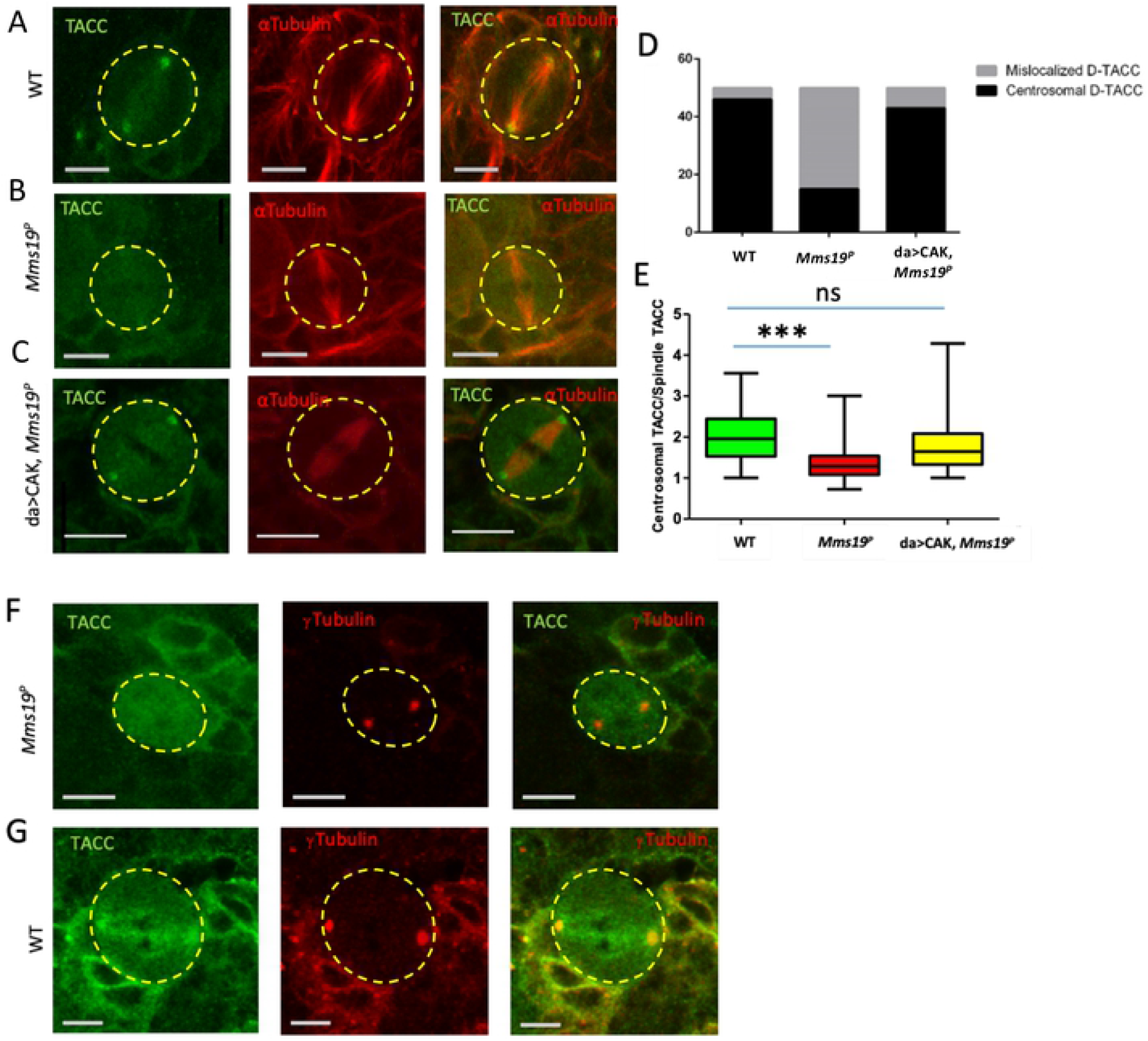
Centrosomal localization of the MT regulator TACC depends on Mms19. (A) A mitotic NB shows TACC localization at the spindle poles. (B, D) >50% of analyzed *Mms19^P^* NBs fail to localize TACC to spindle poles. (C, D) TACC localization was restored upon overexpression of the CAK subunits Cdk7, CycH and Mat1 in the *Mms19^P^* background. (E) The amount of TACC localized on the centrosomes was quantified by comparing the fluorescent intensity of the centrosomal TACC signal to the TACC signal on the spindles. A ratio greater than 1 was considered as clear centrosomal TACC, while a ratio equal to or less than1 indicated unlocalized TACC. SS was calculated by Kruskal-Wallis test, columns were compared by Dunn’s posttest (\*\*\**P*<0.001), scale=5μm. n=60 NBs. (F, G) NBs were stained with antibodies against gTubulin to test for centrosomal localization. (G) TACC co-colocalized with gTubulin on the centrosomes. (F) In the mutant NB TACC fails to concentrate at gTubulin foci.

### Mms19 binds to MTs and stimulates MT assembly

To explore the possibility that Mms19 also functions through different activities, we prepared protein extracts from flies expressing Mms19::eGFP driven by its endogenous promoter (Nag et al, 2018), subjected them to immunoprecipitations and analyzed co-purifying proteins by Mass spectrometry (Supplementary Table S1). Adult wild-type flies and Imp::eGFP expressing flies, respectively, served as controls to exclude any non-specific binding to beads and GFP, respectively. Amongst the proteins exclusively bound to Mms19::eGFP were the CIA proteins Mip18, Ciao1 and Ant2, which form a complex with Mms19 to mediate Fe-S cluster delivery (Gari et al, 2012; Stehling et al, 2012). The fact that we recovered these proteins efficiently, indicated that the purification was efficient. Because *Mms19* functions on microtubules and our second control, IMP::eGFP, is also involved in MT dependent processes, we additionally inspected the data for tubulin and MT binding proteins that are enriched by the Mms19::eGFP compared to the wild-type control without eGFP tag (Supplementary Table S2). This comparison revealed a clear enrichment of tubulin and several Microtubule Associated Proteins (MAPs). Some of the associated proteins were not present at all in the wild-type control, some were present, but at lower levels compared to the Mms19::eGFP and IMP::eGFP fractions. Others were exclusively found in the Mms19::eGFP fraction. These results therefore suggested that Mms19 might directly or indirectly bind to MTs.

To validate the interaction of Mms19 with Tubulin, we next tested whether Mms19::5xHis purified from *E. coli* and α/β-tubulin dimers interact *in vitro*. Mms19::5xHis was incubated at an equimolar ratio with purified porcine brain α/β-tubulin and then bound to the Ni-NTA resin. All incubations and washing steps were carried out at 4°C. Copurifying proteins where then assessed by western blotting. A band corresponding to a-tubulin was observed when tubulin was incubated with Mms19::5xHis (Fig 7A, B), but tubulin alone did not bind to the resin, pointing to a direct interaction between tubulin and Mms19::5xHis.

**Figure 7.**
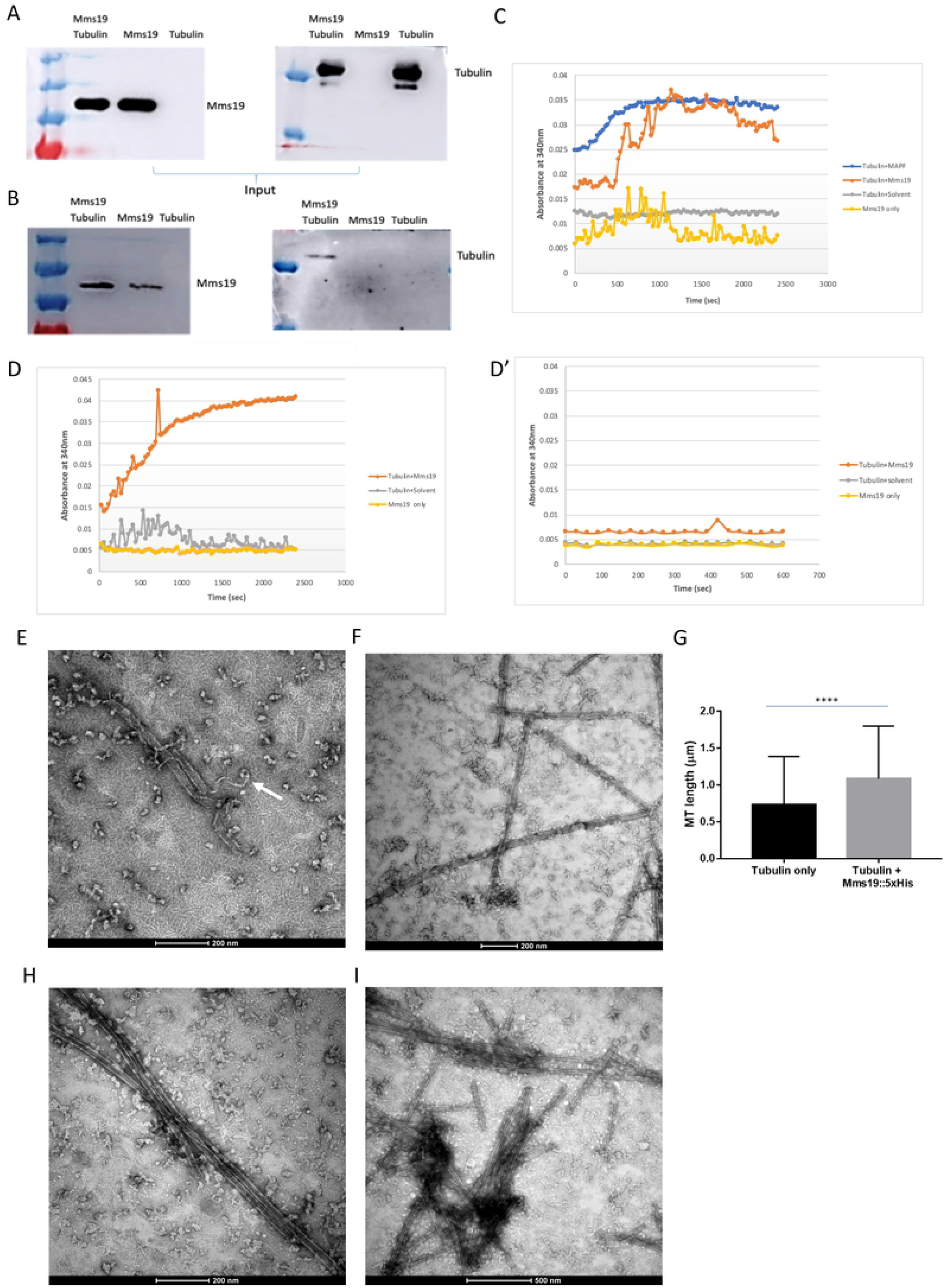
Mms19::5xHis binds to tubulin and stimulates MT assembly *in vitro*. Mms19-Tubulin interaction using a pull-down assay performed at 4°C. Purified tubulin and Mms19::5xHis were first incubated at equimolar ratios and subsequently with Ni-NTA resin which binds His-tags. (A) Input panels show Tubulin added, either alone or with Mms19. (B) Tubulin was found binding to the resin only upon prior incubation with Mms19::5xHis. Tubulin added alone did not bind to the resin. (C) Purified Mms19 was incubated with tubulin in the presence of GTP and the absorbance was measured at 340nm at 35°C to assess polymerization. MAPF (Cytoskeleton Inc.) was used as a positive control, whereas Mms19 alone and tubulin with solvent (the solvent in which Mms19 was reconstituted) were used as negative controls. (D-D’) Absorbance of MTs increases when incubated with Mms19, but sharply decreases after incubating this mixture on ice for 10 minutes. (E) MTs only (without Mms19::5xHis added) visualized by negative stain EM. Here, a MT fiber can be seen depolymerizing (indicated by arrow). (F) In a Mms19::5xHis-MT mixture, discreet particles can be seen binding laterally along the surface of MT. (G) Measuring the length of MT fibers revealed that MTs were generally longer when incubated with Mms19::5xHis and tubulin, compared to tubulin to which only the BRB80/solvent buffer was added. n=150 MTs, ss was calculated using unpaired t-test, ****p<0.0001. MT bundles containing 2 (H) or more than 2 MTs (I) were more frequently observed upon addition of Mms19::5xHis to MTs.

To establish whether Mms19 stimulates MT polymerization, assembly or stability, we performed a spectrophotometric assay in which tubulin was incubated with Mms19::5xHis at a 40:1 ratio (Mirigian et al, 2013). The absorbance of this mixture was recorded at 340nm over 40 min. The increase in absorbance of the sample is a measure for the polymerization of MTs. The positive control, a fraction of known MT elongators (Microtubule Associated Protein Rich Fraction (MAPF) produced a similar maximal absorbance (Fig 7C), but this was not seen when either only Mms19 or only tubulin were added (Fig 7C). Furthermore, to distinguish MT polymerization from aggregation, we incubated this mixture after the polymerization at 4°C. If tubulin indeed polymerizes into MTs, this incubation in the cold should de-polymerize the MTs. On the other hand, if the increasing absorbance was due to aggregation of either tubulin or Mms19, the absorbance due to aggregates would not decrease after cold treatment. Because the absorbance of the sample sharply decreased in the cold, we conclude that Mms19 indeed promoted MT polymerization (Fig 7D-D’). Mms19 thus not only binds to tubulin and MTs, it also stimulates MT polymerization, assembly or stability *in vitro*.

To better understand the mechanism of modulation of the MT dynamics by Mms19, we complemented the biochemical experiments with a high-resolution optical method. Polymerized MTs were incubated at room temperature (RT) with either BRB80/solvent buffer or with Mms19::5xHis, purified upon expression in *E. coli*. These samples were then visualized by negative-stain electron microscopy (EM). MT fibers in general appeared to be less stable upon addition of BRB80/solvent alone (at RT) as some MT fibers were observed undergoing de-polymerization (Fig 7E) under these conditions. Remarkably, MTs incubated with Mms19::5xHis were comparatively longer (Fig 7F, G) and more bundled (Fig 7H, I). Furthermore, the Mms19::5xHis containing samples contained particles not found in the samples without Mms19 and these particles appeared along the lateral MT surface (Fig 7F, H). It thus appears that Mms19 binds along the surface of MTs, stabilizing MTs and possibly stimulating inter-MT contacts.

## Discussion

*Mms19* was initially identified as a gene that regulates the nucleotide excision repair (NER) pathway of DNA repair through its participation in Fe-S cluster assembly. When evidence emerged about a possible NER independent mitotic activity of *Mms19* (Ito et al, 2010; Nag et al, 2018), this warranted further investigation into the precise role of *Mms19* during mitotic spindle assembly in diploid cells. Now we report novel, important roles for *Mms19* during spindle MT assembly and mitotic timing in *Drosophila* NBs. Based on their different suppressibility by elevated CAK levels, *Mms19* phenotypes point to a dual role of the novel mitotic gene *Mms19* in MT assembly. The direct function of Mms19 in binding to, stabilizing and bundling of MTs seems CAK independent and might therefore have an important function in the establishment of other extended microtubule structures, too. Indeed, we found preliminary evidence that this might be the case. Neurites are long projections that emerge in differentiating, postmitotic neurons. They become packed with extended MT bundles which support neurite outgrowth and neuronal signaling. Because these are postmitotc events, this MT formation is not driven by the mitotic kinases. Nevertheless, we found that normal neurite outgrowth depended on Mms19 because *Mms19^P^* neuronal cells were unable to extend neurites in *ex vivo* experiments (Fig S8).

In mitotic NBs, both Mms19 mechanisms, the CAK independent and the CAK dependent, are required. Nag et al. (2018) had presented a model that explained how Mms19 activates the mitotic CAK and the Cdk1 activity. Their model was based on results with larval imaginal discs. We reaffirmed that this *Mms19* activity is also important in NBs, and we further identified a downstream component of this *Mms19*-CAK-Cdk1 mitotic axis, that seems to mediate this activity, the regulator of centrosomal MTs, TACC. In the CAK dependent pathway model (Fig S3A-B), Mms19 competes with the CAK complex for binding to Xpd (Ito et al, 2010; Nag et al, 2018). During mitosis, binding of Mms19 to Xpd prevents the interaction between Xpd and the CAK complex, thus releasing the inhibition of the Cdk-activating kinase activity of CAK by Xpd (Chen et al, 2003; Nag et al, 2018). Our results add an additional dimension to this pathway by linking it to the downstream Aurora A-TACC pathway (Giet et al, 2002; Barros et al, 2005). *Mms19* inactivation prevents normal centrosome localization of TACC and this phenotype is rescued upon expressing additional CAK (Fig 6). Cdk1, the major downstream target of the CAK complex, is known to activate Aurora A kinase during mitosis (Van Horn et al, 2010) and phosphorylation of TACC by Aurora A triggers its centrosomal localization (Giet et al, 2002). Given that the failure of TACC localization in the absence of *Mms19* can be rescued by CAK over-expression, we now propose TACC as a downstream target of the Mms19-CAK-Cdk1 axis and modulation of CAK activity seems to be the main mode of action of Mms19 towards TACC. The best characterized function of TACC is its interaction with Msps which leads to stabilization of astral MTs (Lee et al, 2001). This activity involves their recruitment to the centrosomes, and we found that not only the centrosomal localization of TACC (Fig 6), but also the one of Msps depends on *Mms19* and elevated CAK activity (Fig S6). The activity of *Mms19* through this pathway, which involves the activation of the M-phase regulator Cdk1, therefore contributes significantly to the coordination of cell cycle progression and spindle formation, a process that is affected in the absence of *Mms19* (Fig 2).

Loss of centrosomal TACC or Msps leads to destabilization of centrosomal MTs and defects in chromosome segregation (Gergely et al, 2000; Lee et al, 2001), indicating that the activity of *Mms19* through this pathway makes an important contribution to MT stability and chromosomal segregation. Interestingly, whereas TACC itself may not directly regulate MT assembly, there is good evidence that its partner Msps, which also requires *Mms19* function to localize to centrosomes (Fig S6), can promote MT polymerization. The *Xenopus* homologue of Msps, XMAP215, and the mammalian homolog, chTOG, have been demonstrated to be processive MT polymerases (Brouhard et al, 2008; Gutierrez-Caballero et al, 2015).

CAK overexpression rescued TACC localization, but it could not fully rescue the short spindle/spindle assembly and the microcephaly defects (Figures 1, 3, 5). Furthermore, even if we did not test this directly, we can expect the glial cell MT growth defects (Fig 4B) and the neurite outgrowth defect (Fig S8) to be also independent of the CAK activity because this kinase is mostly active during the mitotic phase of the cell cycle. We found that Mms19 interacts directly with tubulin and MTs. This is interesting because previous work had also suggested an interaction with the mitotic spindle (Ito et al, 2010, van Wietmarschen et al, 2012, Nag et al, 2018). In this study, we report for the first time a direct interaction between Mms19 and tubulin, and we linked this interaction to an activity of Mms19 in stimulating MT assembly (Fig 7). The EM data indicates that Mms19 might contribute to the assembly of MTs by directly binding to them, bundling them and regulating thereby the stability of MTs. Interestingly, such a MT bundling activity seems also crucial for neurite outgrowth, because the initial stage of this process involves bundling of MTs (Miller and Suter, 2018). This would explain why this process is delayed or blocked in most *Mms19^P^* neurons (Fig S8).

Recently, another connection between Fe-S cluster proteins and mitosis was described to involve Kif4a in HEK293 cells (Ben-Shimon et al, 2018). However, its *Drosophila* homologue, Klp3a, was not amongst the proteins co-purifying with Mms19::eGFP in extracts from adult flies (Table S1) and the *Klp3a* phenotypes described by (Williams et al, 1995) are distinctly different from the ones we found for *Mms19^P^*. Nevertheless, it would be interesting to find out whether and where *Mms19* has yet another mitotic function in a metazoan through delivering an Fe-S cluster to Klp3a/Kif4a.

Ito and co-workers first proposed the novel ‘MMXD’ mitotic complex consisting of Mms19, Mip18 and Xpd, that localizes to the mitotic spindle and is required for proper chromosome segregation in human cells (Ito et al, 2010). Interestingly, the epithelial-cell polarity regulator protein Crumbs (Crb) was also shown to form a complex called ‘CGX’ with Xpd and the *Drosophila* homolog of Mip18, Galla-2 (Yeom et al, 2014). Yeom and colleagues found that Crb and Galla-2 associated with the mitotic spindle and were required for spindle assembly and chromosome segregation in young *Drosophila* embryos and that this function could be connected with the modulation of CAK activity by Xpd (Li et al, 2010). Further, a tissue overgrowth phenotype induced by Crb overexpression was rescued by downregulation of Xpd and/or Galla-2. However, the authors did not describe a Crb-Mms19 interaction and Crb did not appear as a Mms19-interacting protein in our Mass-spectrometry screen. Nevertheless, as Mms19 interacts with both Xpd and Galla-2 (Nag et al, 2018), it seems likely that it could be a part of the mitotic ’CGX’ complex. It would thus be interesting to test whether *Mms19* knockdown can also rescue the defects induced by Crb overexpression. Additionally, because neither of the ‘CGX’ proteins are known to have tubulin binding properties, the possibility exists that Mms19 might recruit the CGX/MMXD proteins to the spindles and co-ordinate spindle assembly and chromosome segregation with them.

Multiple MT associated proteins co-operate for the assembly and dynamics of the mitotic spindle. Our *in vitro* studies with *E.coli* expressed Mms19 strongly suggest that Mms19 directly interacts with MTs and that this interaction supports MT growth and bundling. It is possible that *in vivo*, interaction of Mms19 with additional factors might then boost this activity. Correspondingly, in the Mass Spectrometry analysis (Supplementary table 1) we also see MT associated proteins such as Lis-1, Kinesin heavy and light chains. Lis-1 is already known to function during spindle assembly and positioning (Moon et al, 2014). The immunopurification of Mms19 and associated proteins was carried out with extracts from total adult flies, where mitotic cells and particularly cells in M-phase are rare. It is therefore possible that our screen for interactions with Mms19 identified primarily proteins that interact in interphase and that we missed M-phase-specific interactions. Such candidates could be members of the kinesin family, like Kinesin-14 and Kinesin-5, that play important roles during mitosis. For instance, Kinesin-5 drives bipolar spindle assembly by cross-linking and sliding anti-parallel MTs and may also have an additional MT polymerase activity (Fink et al, 2007; Ferenz et al, 2010; Chen and Hancock, 2015; Leary et al. 2019). There is thus an exciting possibility that Mms19 may also bind to other spindle specific MAPs and through its MT modulatory activity serve to assist the function of such MAPs. Further evidence in this direction would reveal invaluable aspects about a possible cross-talk between Mms19 and MAPs.

It was shown recently that MMS19 protein levels are regulated by ubiquitination and subsequent proteasomal degradation in human cells by the ubiquitin ligase MAGE-F1 (Weon et al, 2018). Weon and colleagues showed that the human MMS19 is ubiquitinated at the C-terminal Lysine 993. Alignment of the human and *Drosophila* Mms19 sequences revealed that the human Lys 993 is conserved in *Drosophila* and, additionally, the PTM code 2 software identified this region in *Drosophila* Mms19 as a potential ubiquitinylation site (Minguez et al, 2012, Fig S7). This regulatory mechanism also links MMS19 to certain cancers, especially lung squamous cell carcinomas, where MAGE-F1 was found to be upregulated (Weon et al, 2018). In these tumor cells, up-regulation of MAGE-F1 correlated with increased mutational load and the authors connected these mutations to compromised DNA repair that might lead to chromosomal instability (CIN) due to excessive degradation of MMS19 by MAGE-F1. In support of this view, knock-down of MAGE-F1 considerably slowed tumor growth in a mouse xenograft model. But results from our study as well as previous reports (Ito et al, 2010; Nag et al, 2018) provide strong evidence for the roles of Mms19 in MT assembly and spindle formation and its misfunction in this process could cause CIN, leading to the mutational burden. The mitotic defects we observed upon loss of functional *Mms19* did not lead to high levels of aneuploidy in otherwise healthy NBs (Fig S5). However, in rapidly dividing tumor cells carrying oncogenic driver mutations (and lacking tumor suppressor or mitotic checkpoint functions), spindle defects caused by loss of *MMS19* function could possibly amplify the DNA damage effects and CIN. Indeed, high levels of aneuploidy were observed in human HCT116 cell lines and in *Drosophila* syncytial embryos with reduced *Mms19* function (Ito et al, 2010; Nag et al, 2018). Our results, therefore, provide support for the interpretation that this novel spindle assembly function of *Mms19* might play an important role in preventing CIN and tumor formation in rapidly dividing cells.

Our findings have uncovered novel insights about the mechanism of spindle assembly by *Mms19*. Moreover, as these activities involve a direct association of Mms19 with either Xpd or MTs, the functions described here are most likely independent of the NER role of Mms19. We have also found evidence for a possibly non-mitotic MT regulatory role of *Mms19* for neurite outgrowth. It will be interesting to further analyze the function of Mms19 during the various stages of neurite outgrowth and how its expression and localization is regulated in neurons. A gene that was initially described as a NER regulator, *Mms19* now has the additional roles as a mitotic gene and a MT regulator assigned to it. For the critical roles it fulfils, the proper control of Mms19 expression and localization must be crucial, and erroneous activation of CAK or MT assembly are likely to lead to defects in cell cycle progression. Future studies should therefore address the transcriptional and post-translational control of Mms19 expression and localization and how this impacts its functions in cell physiology, development and diseases.

## Materials and Methods

### Key resources tables

#### The following table lists the fly stocks used in this study

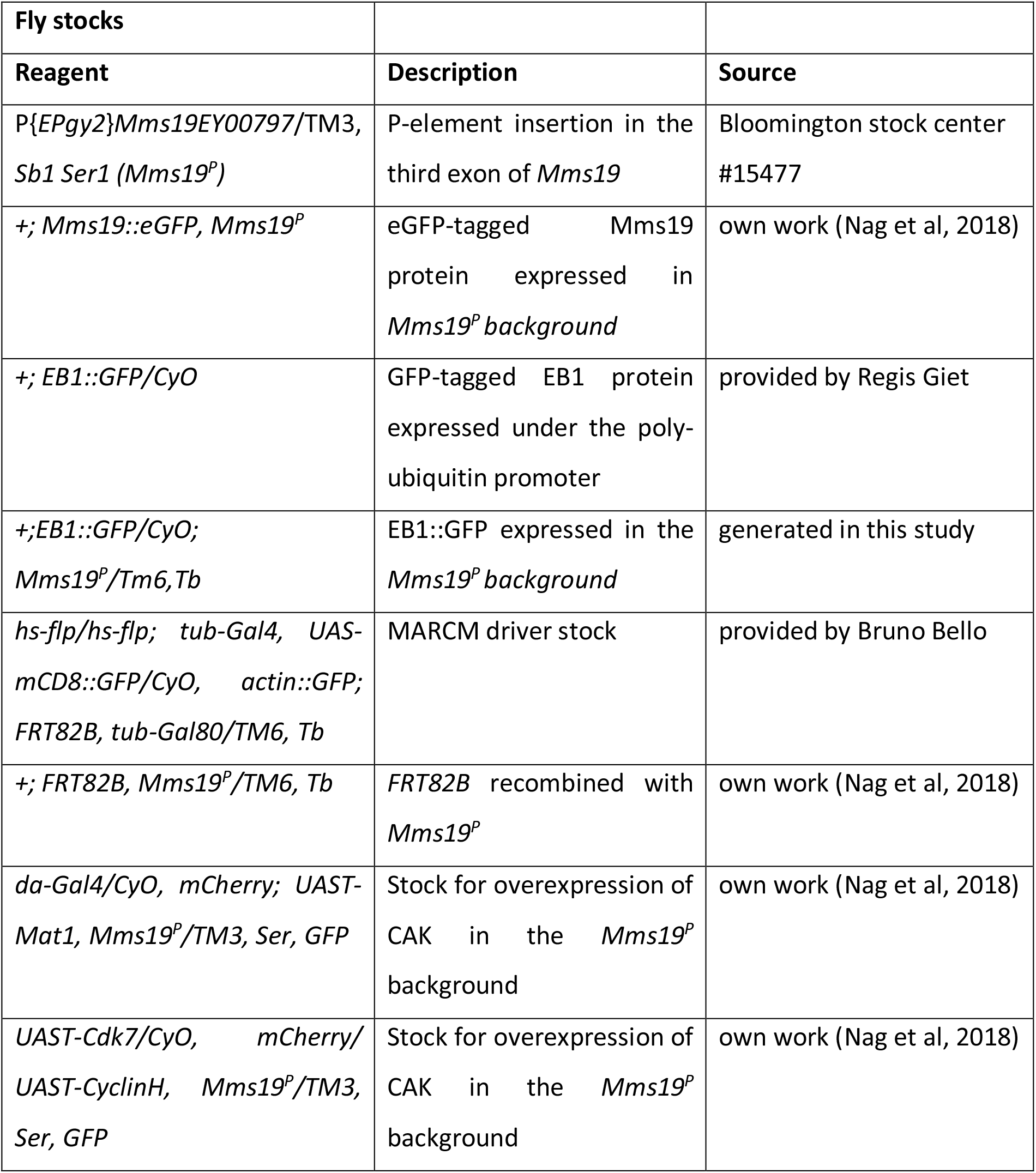

| Fly stocks |  |  |
| --- | --- | --- |
| Reagent | Description | Source |
| P{EPgy2}Mms19EY00797/TM3, Sb1 Ser1 (Mms19<sup>P</sup>) | P-element insertion in the third exon of *Mms19* | Bloomington stock center #15477 |
| +; Mms19::eGFP, Mms19<sup>P</sup> | eGFP-tagged Mms19 protein expressed in *Mms19<sup>P</sup>* background | own work (Nag et al, 2018) |
| +; EB1::GFP/CyO | GFP-tagged EB1 protein expressed under the poly-ubiquitin promoter | provided by Regis Giet |
| +; EB1::GFP/CyO; Mms19<sup>P</sup>/Tm6, Tb | EB1::GFP expressed in the *Mms19<sup>P</sup>* background | generated in this study |
| hs-flp/hs-flp; tub-Gal4, UAS-mCD8::GFP/CyO, actin::GFP; FRT82B, tub-Gal80/TM6, Tb | MARCM driver stock | provided by Bruno Bello |
| +; FRT82B, Mms19<sup>P</sup>/TM6, Tb | FRT82B recombined with *Mms19<sup>P</sup>* | own work (Nag et al, 2018) |
| da-Gal4/CyO, mCherry; UAST-Mat1, Mms19<sup>P</sup>/TM3, Ser, GFP | Stock for overexpression of CAK in the *Mms19<sup>P</sup>* background | own work (Nag et al, 2018) |
| UAST-Cdk7/CyO, mCherry/ UAST-CyclinH, Mms19<sup>P</sup>/TM3, Ser, GFP | Stock for overexpression of CAK in the *Mms19<sup>P</sup>* background | own work (Nag et al, 2018) |

#### The following table lists the primary and secondary antibodies used for immunostainings

The following table lists the primary and secondary antibodies used for immunostainings:
| **Primary antibodies** |  |  |  |
| --- | --- | --- | --- |
| **Antibody** | **Manufacturer** | **Catalog number** | **Dilution** |
| Rat anti-Miranda | Abcam | Ab197788 | 1:250 |
| Rabbit anti-alpha Tubulin | Abcam | Ab18251 | 1:500 |
| Mouse anti-alpha Tubulin | Sigma | T6199 | 1:500 |
| Rabbit anti-GFP | Immunokontakt | 210-PS-1GFP | 1:500 |
| Rabbit anti-pH3 | Cell Signaling | 9701 | 1:200 |
| Rabbit anti-Mms19 | Genscript | Custom antibody | 1:500 |
| Mouse anti-γH2av | DSHB | UNC93-5.2.1 | 1:20 |
| Rabbit anti-PKC $\zeta$ (aPKC) | Santa-Cruz<br>Biotechnology | sc-216 | 1:2,000 |
| Mouse anti-Futsch | DSHB | 22C10 | 1:20 |
| Mouse anti-γTubulin | Sigma | T6557 | 1:1,000 |
| Rabbit anti-TACC | provided by Jordan Raff | - | 1:1,000 |
| Rat anti-Elav | DSHB | 7E8A10 | 1:20 |
| Rabbit anti-Msps | Provided by Hiro<br>Ohkura | - | 1:1,000 |
| **Secondary antibodies** |  |  |  |
| Goat anti-rat Cy3 | Jackson Immuno | 112-165-167 | 1:150 |
| Goat anti-mouse Alexa 488 | Invitrogen | R37120 | 1:500 |
| Goat anti-rabbit Alexa 488 | Invitrogen | A27034 | 1:500 |
| Goat anti-mouse Alexa 647 | Invitrogen | A21235 | 1:500 |

#### The following table lists the antibodies used for probing western blots

The following table lists the antibodies used for probing western blots:
| Primary antibodies |  |  |  |
| --- | --- | --- | --- |
| Antibody | Manufacturer | Catalog number | Dilution |
| Rabbit anti-Mms19 | Synthesized by<br>Genscript Inc. | Custom antibody | 1:2,000 |
| Rabbit anti-alpha<br>Tubulin | Abcam | Ab18251 | 1:2,000 |
| Secondary antibodies |  |  |  |
| Goat anti-rabbit HRP | Thermo Fischer<br>Scientific | 65-6120 | 1:10,000 |

#### Following table lists the kits and reagents used in this study

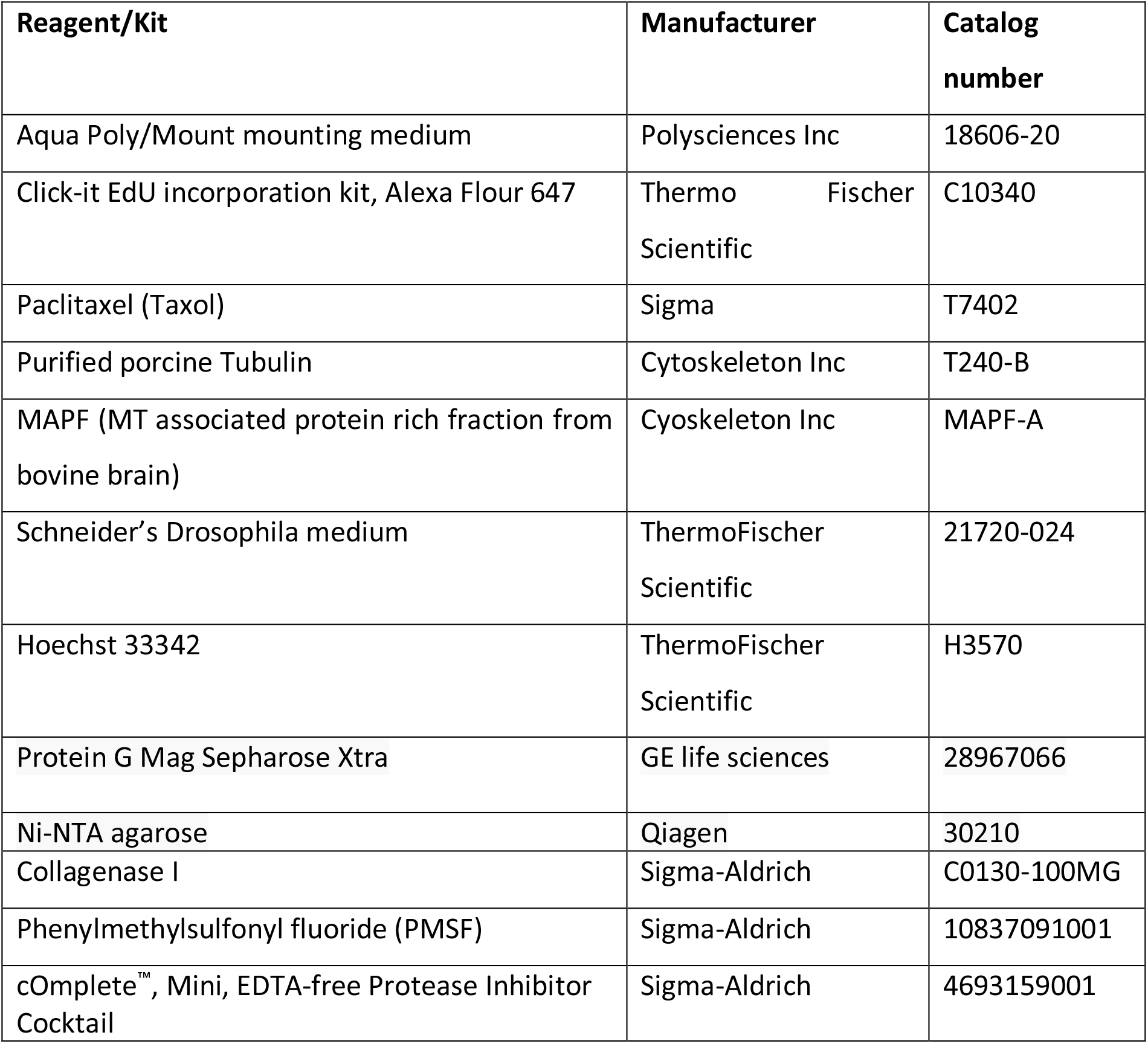
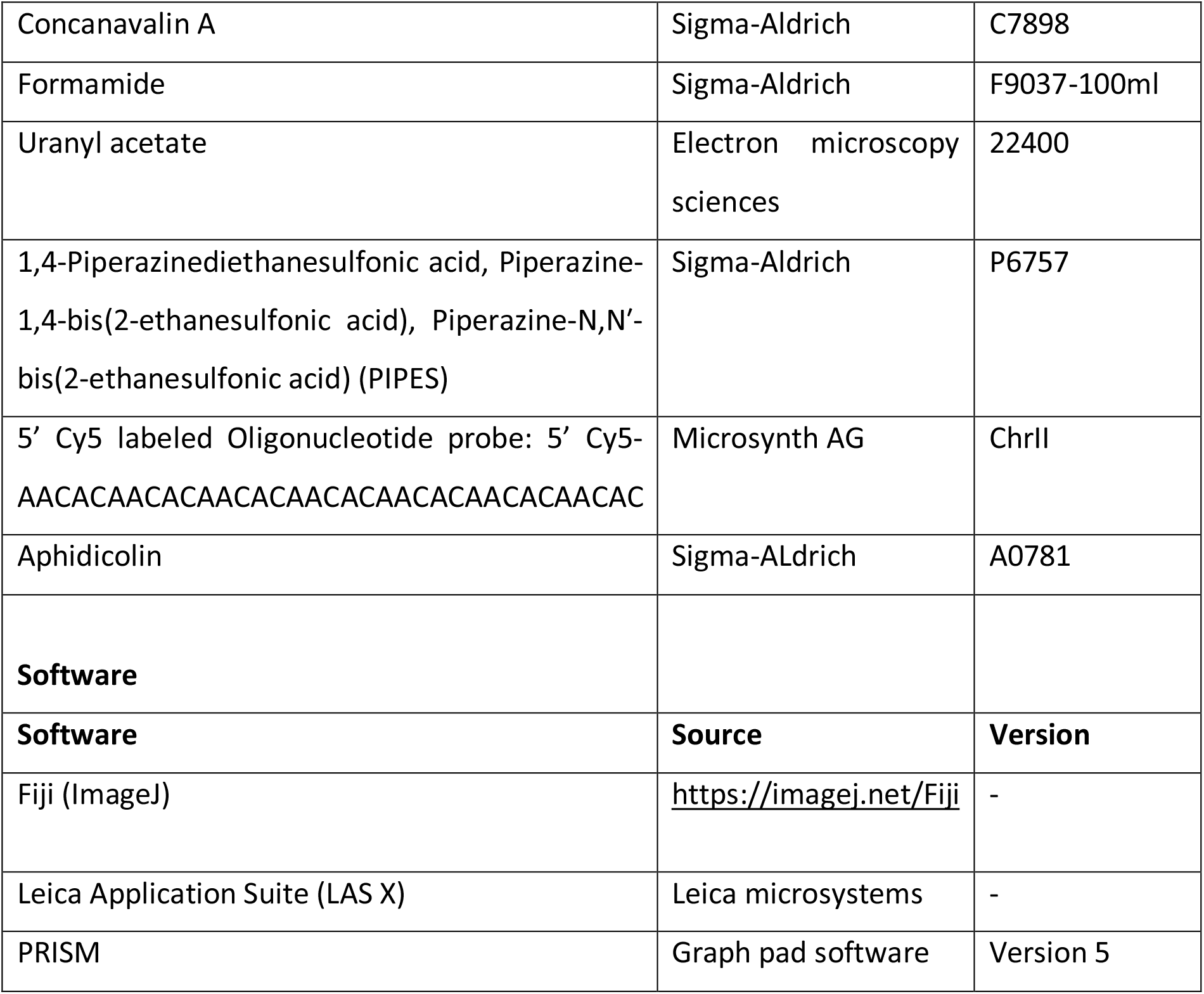

## Method Details

### Dissection and immunostaining of larval brains

Larval brains were dissected and stained as described (Daul et al, 2010). Briefly, wandering third instar larvae were dissected in PBS and fixed for 15min in 4% paraformaldehyde supplemented with 5mM MgCl2 and 1mM EGTA (to stabilize MTs). After 3 washes in PBS+0.1% TritonX-100 (PBST), primary antibodies were added. After overnight incubation, the brains were washed 3x with PBST, stained with secondary antibodies and mounted on glass slides in Aqua Polymount mounting medium (Polysciences).

For MT regrowth assays, brains were dissected in Schneider’s medium (supplemented with 10% fetal bovine serum) at 25°C and incubated on ice for 30 min to depolymerize MTs. Brains were then incubated for different time points in a 25°C water bath, followed by fixation and immunostaining. This experiment was performed three times and 20 spindles were analyzed in each iteration. Slides were imaged on a Leica TCS-SP8 microscope (Leica Microsystems) equipped with a 63X, NA 1.4 Plan Apochromat objective. Images were acquired using LAS X software and analyzed using Fiji/ImageJ (Schindelin et al., 2012).

### Measuring brain compartment volumes

Whole brain lobes were immunostained with antibodies against Miranda (Mira) and phospho-Histone 3 (pH3), and DNA was visualized with Hoechst 33342. Z stacks were acquired with a TCS-SP8 confocal microscope on a 63X, 1.4NA Plan-Apochromat objective with 1μm spacing between each optical section. Segmentation, volume measurement and 3D reconstruction was performed by using the TrackEM2 plugin in Fiji (Cardona et al, 2012).

### MARCM crosses

For generating mosaic clones the driver stock *hs-flp; tub-Gal4, UAS-mCD8::GFP/CyO, actin::GFP; FRT82B, tub-Gal80/TM6*, Tb was crossed to *+; FRT82B, Mms19^P^/TM6, Tb*. GFP-Balancer negative 24hr old larvae were selected and heat shocked at 37°C in a glass vial submerged in a water bath for 15min. Larvae were then returned to 25°C and the brains of the non-Tubby larvae were dissected 48hrs later.

### Live imaging

Brains expressing EB1::GFP were dissected and mounted on stainless steel chambers as described in (Cabernard et al, 2013). Brains were then imaged using a 100X, NA 1.3 oil immersion objective on a Visiscope Spinning disk microscope (Visitron GmbH) fitted with Nikon Ni2 stand and a Photometrics Evolve 512 EMCCD camera. Images were acquired for 60 seconds at of 500ms time intervals at 200ms exposure at 60% laser power (488nm). To track the particle velocities, the particles were manually traced in Fiji/ImageJ (Schindelin et al., 2012). Probably due to limited resolution, it was not possible to unambiguously account for merging or splitting events and therefore only particles with a linear trajectory, which did neither split nor merge, were analyzed. At least 5 particles from each spindle were analyzed from a total of 11 cells per genotype. For surface glia quantification, at least 10 particles were analyzed from each of the 15 brains. Stacks were exported to .AVI movies at 14 frames per second.

To measure the duration of mitosis, EB1::GFP expressing brains were mounted as described above and imaged at 40% laser power with a 63X, 1.3 NA objective of a Nikon W1 LIPSI spinning disk microscope fitted with a Photometrics Prime 95B CMOS camera. Movies were acquired on Nikon’s NIS elements software for 2hrs with an interval of 1 min at 200 ms exposure. Z-stacks were acquired simultaneously with 2μm distance between successive optical sections. Movies were analyzed and processed with Fiji/ImageJ. Stacks were exported to .AVI movies at 12 frames per second.

### Quantification of spindle orientation

The orientation of the mitotic spindle was examined with respect to the basal Mira crescent. A reference line was drawn passing approximately through the center of the Mira crescent and the angle between the spindle and this reference line was determined using the ImageJ angle tool.

### EdU incorporation

Click-it EdU kit (Invitrogen) was used to measure EdU incorporation. Brains were dissected in PBS and incubated in Schneider’s medium supplemented with 10μM 5-Ethynyl-2’-deoxyuridine (EdU) for 2hrs at 25°C. EdU is an analogue of Thymidine and is incorporated by S phase cells during DNA replication. EdU is then detected due to its binding to a dye-azide conjugate. Brains were subsequently fixed with 4% PFA and incubated with primary antibodies (anti-Mira to mark NBs and anti-pH3 to mark mitotic cells) and secondary antibodies. Subsequently, they were processed for EdU detection following the manufacturer instructions.

### Preparation of whole-fly extract

1g of flies were collected in Eppendorf tubes and frozen in liquid nitrogen. Frozen flies were crushed into a fine powder using a pre-cooled mortar pestle. The powder was incubated in lysis buffer (25mM Hepes, 150mM NaCl, 1mM EDTA, 0.1% TritonX-100, 1mM phenylmethylsulfonyl fluoride (PMSF), 1 complete EDTA-free protease inhibitor tablet (Roche/Sigma-Aldrich)) for 30 min and then centrifuged in an Eppendorf tube for 30 min at 16,000 g and 4°C. The supernatant was saved and snap frozen in liquid nitrogen.

### Immunoprecipitation to prepare extract for Mass Spectrometry

ProteinG-Mag Sepharose (GE) beads were washed 3X with PBS, incubated for 2hrs with anti-GFP antibody. Beads were then incubated with the crude extracts for 6hrs at 4°C. Following 3 washes with the wash buffer (25mM Hepes, 150mM Nacl, 1mM EDTA, 1mM PMSF, 1 tablet complete EDTA free protease inhibitor tablet), bound proteins were eluted by 15 min incubation in urea elution buffer (6–8 M Urea, 20 mM Tris pH 7.5, and 100 mM NaCl) or glycine elution buffer (100mM Glycine, pH 2.6. These eluates were neutralized by adding 150mM Tris-Cl, p 8.8).

### Mass-Spectrometry

Eluted proteins in 8M urea were processed essentially as described by Engel and colleagues (Engel et al, 2014). Briefly, proteins were reduced by addition of 1/10 volume of 0.1 M DTT and incubated for 30 min at 37°C, followed by alkylation with a five-fold molar excess of iodoacetamide and incubation for 30 min at 37°C. Proteins were precipitated at −20°C by addition of 5 volumes cold acetone and incubation for 30 min at −20°C. All liquid was carefully removed, and the pellet dried in ambient air for 15 min before reconstitution of the proteins in 8 M urea, 50 mM Tris-HCl pH 8.0 to a final protein concentration between 0.2-0.3 mg/mL. Protein concentration was determined by Bradford assay. An aliquot corresponding to 5 μg protein was diluted to a final urea concentration of 2 M urea with 20 mM Tris-HCl pH 8.0, and 2 mM CaCl2. Proteins were digested by trypsin (1:50 (w/w) trypsin/protein ratio) for 6 hours at 37°C. The digests were acidified with TFA (1%) and analyzed by LC-MS/MS (EASY-nLC 1000 coupled to a QExactive HF mass spectrometer, ThermoFisher Scientific) with three repetitions injecting an aliquot of 500 ng protein. Peptides were trapped on an Acclaim PepMap100 C18 pre-column (3μm, 100 Å, 75μm x 2 cm, ThermoFisher Scientific, Reinach, Switzerland) and separated by backflush on a C18 column (3μm, 100 Å, 75μm x 15 cm, Nikkyo Technos, Tokyo, Japan) by applying a 40 min gradient of 5% acetonitrile to 40% in water, 0.1% formic acid, at a flow rate of 300 nl/min. Peptides of m/z 400-1400 were detected at a resolution of 60,000 m/z 250 with an automatic gain control (AGC) target of 1E06 and maximum ion injection time of 50 ms. A top fifteen data dependent method for precursor ion fragmentation was applied with the following settings: resolution 15,000, AGC of 1E05, maximum ion time of 110 ms, charge inclusion of 2+ to 7+ ions, peptide match on, and dynamic exclusion for 20 sec, respectively.

Fragment spectra data were converted to mgf with ProteomeDiscoverer 2.0 and peptide identification made with EasyProt software searching against the forward and reversed UniprotKB *Drosophila melanogaster* protein database (Release 2016_11), complemented with commonly found protein sequences of contaminating proteins, with the following parameters: parent mass error tolerance of 10 p.p.m., trypsin cleavage mode with three missed cleavages, static carbamidomethylation on Cys, variable oxidation on Met and acetylation on protein N-terminus. On the basis of reversed database peptide spectrum matches, a 1% false discovery rate was set for acceptance of target database matches, and only proteins with at least two different peptide sequences identified were allowed.

### Immunoprecipitation/Pull-down assays

50μg of purified Mms19 (Tagged with 5X Histidine at C-terminus, synthesized by Genscript Inc and solubilized in 20mM Tris, 150mM NaCl, 0.5 M Arginine) was incubated with 50μg purified porcine tubulin (Purchased from Cytoskeleton Inc) at 4°C for 2 hrs. This mixture was subsequently incubated with Ni-NTA agarose (equilibrated with 20mM Tris-Cl and 250mM NaCl) for 1 hr at 4°C. The beads were then washed three times with wash buffer containing 20mM Tris, 250mM NaCl and 20mM Imidazole and the bound proteins were eluted by adding elution buffer (20mM Tris, 250mM NaCl and 500mM Imidazole) to the resin on ice for 10 min. The eluate was then analyzed by probing Western blots with anti-Mms19 antibodies and rabbit anti-alpha tubulin antibodies.

### *In vitro* MT turbidity assay

40μM solution of tubulin (porcine brain, Cytoskeleton, Inc) was prepared in Brinkley’s buffer (BRB80, which contains 80mM PIPES buffer, 1mM EGTA, 2mM MgCl2) +10% glycerol and supplemented with 1mM GTP. The tubulin solution was then incubated with test proteins (MAPF 1mg/ml, Mms19 0.16mg/ml (concentration of Mms19 used was 1μM)) at 37°C and absorbance was measured at 340nm on a SpectraMax 340PC microplate reader (Molecular devices). Two negative controls were used: Mms19 only and Tubulin incubated with the solvent in which Mms19 is solubilized (20mM Tris-CL, 150mM NaCl, 0.5M Arginine). Results were validated from four independent experiments.

### Neuronal *in vitro* cultures

Dissociated brain cells were cultured *in vitro* according to a protocol previously described (Egger et al, 2013). Briefly, 15-20 third instar larvae were dissected in PBS, and washed 3 times with Rinaldini solution (800 mg NaCl, 20 mg KCl, 5 mg NaH2PO4, 100 mg NaHCO3, 100 mg glucose, in 100 ml distilled water). The brains were then incubated in 0.5% collagenase I in Rinaldini solution for 60 min and subsequently washed 4 times in Schneider’s medium. The treated tissues were then dissociated by pipetting 100-200 times. The resulting cell mixture was passed through a 40μm mesh to remove cell clusters, and the brain cells were then incubated for 24hrs at 25°C. After 24hrs incubation, the cells were plated on concanavalin A-coated coverslips, fixed and stained.

### Preparation of MTs for EM

To obtain polymerized MTs, 20μM tubulin was incubated in the presence of 10μM taxol at 37°C for 30min. The sample was then subjected to ultra-centrifugation for 10min at 100,000 g at 4°C in a Beckman Airfuge ultracentrifuge. The supernatant containing un-polymerized dimers was then removed and the pellet was reconstituted with either 15μl BRB80/solvent (BRB80 components: 80mM PIPES buffer, 1mM EGTA, 2mM MgCl2 + Mms19 solvent constituents: 20mM Tris-CL, 150mM NaCl, 0.5M Arginine) or 15 μl of 0.5μM Mms19::5xHis in BRB80/solvent.

### Negative stain EM

5 μl of samples were applied on glow discharged, carbon-coated copper EM grids for 1min. The excess sample was then washed off by dipping the grid in milli-Q water. The sample on the grid was then fixed/negatively stained with 2% Uranyl acetate for 30 sec and then the excess fluid was removed by using filter paper. The samples were imaged at a nominal magnification of 63,000x or 87,000x on a FEI Tecnai Spirit EM operated at 80eV and fitted with a digital camera. MT length and number of MT bundles were quantified from images obtained from three independent experiments.

### Fluorescent *in situ* hybridization (FISH)

The protocol for FISH was adapted from Dernburg, 2011 with minor modifications. Briefly, third instar larval brains were dissected in PBS and fixed with 4% PFA. The brains were then washed 3 times for 10min each with 2xSSCT (0.3M Sodium Chloride, 0.03M Sodium Citrate and 0.1% Tween 20) followed by 10min washes respectively with 2xSSCT/20% Formamide, 2xSSCT/40% Formamide, 2xSSCT/50% Formamide. 100ng of the oligonucleotide probe: 5’ Cy5-AACACAACACAACACAACACAACACAACACAACAC that binds to a specific region on the 2^nd^ chromosome (Dernburg, 2011) was then added to the Hybridization buffer (20% dextran sulfate, 2xSSCT, 50% Formamide) and this solution was incubated with the brains in a PCR tube. The probes were then denatured at 92°C for 3 min and then allowed to anneal with the chromosomal DNA overnight at 37°C. The sample was then washed thrice at 37°C with the following solutions for 20min each: 2xSSCT/50% Formamide, 2xSSCT/40% Formamide, 2xSSCT/20% Formamide. After two more washes with 2xSSCT, the sample was stained with Hoechst, mounted using Aqua-poly mount, and imaged on a Leica SP8 confocal microscope with a 63x objective.

### Statistical analysis

Data was analyzed using Graph Pad prism 5.0 software. Means of two groups were compared and significance calculated by using unpaired students t-test. Multiple groups were compared, and significance calculated using the Kruskal-Wallis test and the Dunn’s post-test.

## Acknowledgments

We would like to thank Alex Bird, Carlo Largiader and our group members for helpful discussions and feedback. Our thanks also go to Regis Giet, Bruno Bello, Claudio Sunkel, Hiro Ohkura, Anne Marcil, Jordan Raff, the Bloomington Stock Center (University of Indiana), DSHB (University of Iowa) for providing antibodies and fly stocks, and FlyBase for excellent community support. The authors also wish to thank Manfred Heller, Sophie Braga and the Proteomics Mass Spectrometry Core facility at the University of Bern, Yury Belyaev of the Microscopy Imaging Center (University of Bern), and Beat Haenni and Benoit Zuber for their services and support.

## Supporting Information legends

### Supporting Figures

**Figure S1: Higher fraction of Mms19P NBs in mitosis.** (A) NBs were classified into 1) G1/G0 phase if they did not stain for either EdU or pH3; 2) S phase if the NBs stained positively for EdU; 3) M phase for NBs staining positively only for pH3 and 4) G2/M for cells staining positively for both EdU and pH3. The G2/M cells are cells transitioning from G2 to M phase. NBs in each phase were counted per brain lobe and this data was represented as percentage (e.g. if in one brain lobe 20 out of 100 NBs were pH3 positive, then 20% cells were classified as in M phase). The percentages for each phase were compiled and compared per brain lobe across the 4 genotypes. Box plot charts represent percentage of cells in (B) G1/G0 phase, (C) S phase and (D) G2/M phase. (E) M phase. 30 brain lobes per genotype were analyzed. SS was calculated using Kruskal-Wallis test, columns compared using Dunn’s post test, ***(*P*<0.001), *(*P*<0.05). Scale=5μm.

**Figure S2: *Mms19* is cell autonomously required to maintain normal cell numbers in MARCM clones.** (A)-(C) In order to study cell cycle progression in a single NB lineage, we used the Mosaic Analysis with a Repressible Cell Marker (MARCM) technique (Lee and Luo, 2001). This technique utilizes the UAS-GAL4-GAL80 system and the FLP-FRT recombination system. With this technique, a population of cells arising from the same progenitor can be specifically labeled. Additionally, the progenitor cell can carry a mutation along with a GFP marker. Defects in this cell, along with its progeny can be analyzed in an otherwise WT background. (E) MARCM clones were induced in NBs in 24hrs old larvae. These larvae were dissected after another 48hrs to determine the number of cells per clone in *Mms19P* and WT/control clones. (F) The graph shows a significant reduction in numbers of cells in mutant clones. SS was determined by an unpaired t-test (\*\**P*<0.01), scale=5μm, n=60 clones from each genotype.

**Figure S3: Model for mitotic and post-mitotic pathways of Mms19. (**A) Mms19 binds to Xpd, thereby releasing the CAK complex to activate its mitotic targets. (B) Downregulation of Mms19 by mutations, knock-down or proteasome mediated degradation allows Xpd to associate with CAK and core TFIIH, thereby targeting Cdk7 activity away from the mitotic targets and towards transcriptional targets like the PolII-CTD (Cameroni et al, 2010). (C) Mms19 binds to MTs and promotes MT stability and bundling, contributing significantly to establishing extended MT structures such as mitotic spindles and possibly the MTs driving neurite outgrowth.

**Figure S4: *Mms19P* NBs show spindle orientation defects.** (A) Spindle orientation in the WT NBs is tightly coupled to the apical-basal polarity axis. (B) In a fraction of *Mms19P* NBs the spindle appears mis-oriented with respect to the Mira crescent localization. (D-E) Quantification of the spindle angle revealed that a considerably higher number of *Mms19P* NBs are oriented at an angle of more than 10° w.r.t Mira. n=65, scale=5μm.

**Figure S5: *Mms19P* NBs do not display high levels of aneuploidy.** (A) WT, *Mms19P* and da>CAK, *Mms19P* brains were fixed and fluorescent *in situ* hybridization was performed on them. Cy5-labelled DNA probes that specifically bind to regions on the 2nd chromosome were used to determine the number of chromosomes. (A) the signal is seen as 2 dots in the WT NBs corresponding to the diploid state of the cell. (B) An example of an aneuploid *Mms19P* NB showing 3 dots. (C) However, aneuploid cells in both *Mms19P* and da>CAK, *Mms19P* NBs were extremely rare. SS was calculated using Fisher’s exact test.

**Figure S6: *Mms19* is necessary for centrosomal localization of Msps in NBs.** (A, C) WT and da>CAK, *Mms19P* NBs showing Msps localization on centrosomes and spindles. (B) In *Mms19P* NBs, Msps does not concentrate on centrosomes. (D) To quantify centrosomal accumulation of Msps, an analysis similar to that done in Fig 6 was performed. SS was calculated using Kruskal-Wallis test, columns compared using Dunn’s post test, ***(*P*<0.001), scale=5μm, n=30.

**Figure S7:** Putative ubiquitination site in *Drosophila* Mms19: A pairwise sequence alignment of the human MMS19 and *Drosophila* Mms19. The alignment performed was based on the Needleman-Wunsch algorithm (Needleman and Wunsch, 1970; https://www.ebi.ac.uk/Tools/psa/emboss_needle/). Analysis of both sequences by the PTMcode 2 software indicates that the C-terminal K993 site in the Human protein is conserved in the *Drosophila* Mms19 at K921.

**Figure S8: *Mms19P* neurons are unable to form long neurites in culture.** (A) WT NBs differentiated into neurons and extended neurites after 24hrs in culture. Neurons were positive for Elav (marker for post mitotic neuronal nuclei). (B) *Mms19P* neurons stained positively for Elav after 24hrs in culture, indicating that these neurons had differentiated. (C) *Mms19P* neurons, did not grow extended neurites after 24hrs or even (D) 48hrs in culture. Scale bar=5μm.

### Supporting Tables

**Supplementary Table 1.** The table lists proteins found to exclusively co-purify with Mms19::eGFP. CIA proteins, which are already known to form a complex with Mms19, are highlighted in Red. Microtubule associated proteins are highlighted in Green.

**Supplementary Table 2.** Proteins that bound to the anti-GFP antibody coated beads from all three fly extracts are listed. These include different isoforms of α-and βtubulin that were identified in all extracts. However, the PMSS scores are higher for the tubulin isoforms immuno-precipitating from the Mms19::eGFP extract. This indicates that tubulin was enriched in the Mms19::eGFP fraction compared to the wild-type control.

### Supporting Movies

**Mov 01:**

**Analysis of mitosis duration in WT NB:** WT brains expressing EB1::GFP were dissected, mounted on a stainless steel chamber (Cabernard et al, 2013) and NB mitosis was imaged with a 63x objective on a spinning disk confocal microscope. Mitosis duration was measured from NEBD onset (start at 0 min) until cytokinesis. Scale=5μm

**Mov 02:**

**Analysis of mitosis duration in *Mms19^P^* NB:** Mitosis was visualized in *Mms19^P^* brain NBs expressing EB1::GFP. On average, *Mms19^P^* brain NBs took twice as long as WT NBs to finish mitosis. In this presented case, the NB completes mitosis 22min after NEBD onset with cytokinesis. Scale=5μm

**Mov 03:**

**Spindle assembly defect in *Mms19^P^* NB:** In around 10% of EB1::GFP expressing *Mms19^P^* brain NBs, the spindle starts assembling before the centrosomes have migrated to the opposite sides, forming a ‘kinked’ spindle at 4min post NEBD onset. This indicates that Mms19 is needed for coordinating spindle formation with cell cycle progression. Scale=5μm

**Mov 04:**

**Analysis of EB1::GFP labelled MT velocity in WT NBs**: In order to determine the speed of growing MT tips, WT brains expressing EB1::GFP were dissected, mounted on a stainless steel chamber (Cabernard et al, 2013) and spindles were imaged with a 100x objective on a spinning disk confocal microscope. Images were acquired at an interval of 500ms for 1min. Scale=5μm.

**Mov 05:**

**Analysis of EB1::GFP labelled MT velocity in *Mms19^P^* NBs:** Live imaging was performed on *Mms19^P^* brain NB spindles to determine the velocity of growing MT tips. Images were acquired at an interval of 500ms for 1min. The spindle shown here appears to tilt repeatedly, perhaps due to defects in astral MT assembly (Fig 2, Fig 6). Scale=5μm.

**Mov 06:**

**Measuring MT plus tip speeds in post mitotic surface glia of WT brains:** In order to determine whether *Mms19* affects MT growth in post mitotic cells, WT brains expressing EB1::GFP were dissected, mounted on a stainless steel chamber (Cabernard et al, 2013) and MT growth in surface glia was imaged with a 100x objective on a spinning disk confocal microscope. Images were acquired at an interval of 500ms. Scale=5μm.

**Mov 07:**

**Measuring MT plus tip speeds in post mitotic surface glia of *Mms19^P^* brains:** MT growth in EB1::GFP expressing *Mms19^P^* surface glia was visualized using spinning disk confocal microscopy. Images were acquired at an interval of 500ms. Scale=5μm.

